# Overlap of expression and alignment of diurnal and circadian rhythmicity in the human blood transcriptome with organ and tissue specific rhythmicity in a non-human primate

**DOI:** 10.1101/2021.08.13.456271

**Authors:** Carla S Möller-Levet, Emma E Laing, Simon N Archer, Derk-Jan Dijk

## Abstract

**BACKGROUND:** Twenty-four-hour rhythmicity in transcriptomes of tissues and organs is driven by local circadian oscillators, systemic factors, the central circadian pacemaker, and light-dark cycles. This rhythmicity is to some extent organ- and tissue-specific such that the sets of rhythmic transcripts or their timing are different across tissues/organs. Monitoring rhythmicity of tissues and organs holds promise for circadian medicine, but in humans most tissues and organs are not easily accessible. To investigate the extent to which rhythmicity in the human blood transcriptome reflects rhythmicity in tissues and organs, we compared the overlap and timing of rhythmic transcripts in human blood and rhythmic transcripts in 64 tissues/organs of the baboon.

**METHODS:** Rhythmicity in the transcriptomes of humans and baboons were compared using set logic, circular cross-correlation, circular clustering, functional enrichment analyses and partial least squares regression.

**RESULTS:** Of the 759 orthologous genes that were rhythmic in human blood, 652 (86%) were also rhythmic in at least one baboon tissue. Most of these genes were associated with basic processes such as transcription and protein homeostasis. 109 (17%) of the 652 overlapping rhythmic genes were reported as rhythmic in only one baboon tissue or organ and several of these genes have tissue/organ-specific functions. Analysis of the alignment between baboon and human transcriptomes showed that in these diurnal species, rhythmicity is aligned with the onset, rather than midpoint or end of light period. In both species, the timing of rhythmic transcripts displayed prominent ‘night’ and ‘day’ clusters, with genes in the dark cluster associated with translation. The timing of human and baboon transcriptomes was significantly correlated in 25 tissue/organs with an average earlier timing of 3.21 h (SD 2.47 h) in human blood.

**CONCLUSIONS:** The human blood transcriptome contains sets of rhythmic genes that overlap with rhythmic genes of tissues/organs, some of which are tissue/organ-specific, in the baboon. The rhythmic sets vary across tissues/organs but the timing of most rhythmic genes is similar across human blood and baboon tissues/organs. These results have implications for our understanding of the regulation of rhythmicity across tissues/organs and species and development of blood transcriptome-based biomarkers for rhythmicity in tissues and organs.

## Background

Twenty-four-hour rhythmicity is a prominent characteristic of behaviour, physiology and molecular processes in single- and complex multi-cellular eukaryotic organisms including humans and other primates [1, 2]. When this rhythmicity is observed in the presence of 24-h environmental cycles it is referred to as diurnal rhythmicity. Near 24-h rhythmicity that persists in the absence of environmental cycles qualifies as circadian rhythmicity [2]. Circadian rhythmicity is generated by a molecular ‘clock’ comprising a transcription/translation feedback loop consisting of core clock genes such as Arntl, Clock, Cry1, Cry2, Per1, Per2, Per3. This core clock drives circadian rhythmicity in gene/protein expression present in nearly all organs, tissues, and cell types [3].

In mammals, rhythmicity in peripheral tissues and organs is synchronised to the 24-h day by neural and endocrine rhythms, such as rhythmicity in the autonomic nervous system and the cortisol rhythm driven by the suprachiasmatic nulcei (SCN), which in turn is synchronised to the 24-h light-dark cycle via neural connections from the retina and its photoreceptors [4]. Tissue- and organ-specific temporal programs are, however, also affected by behavioural and associated systemic factors. For example, in mice, rhythmicity in the liver transcriptome is driven by ‘clock genes’ as well as feeding [5]. Likewise, in the mouse brain, rhythmicity is largely driven by the sleep-wake cycle and to a lesser extent by the circadian molecular clock [6].

In humans, twenty-four-hour rhythmicity in the brain is disrupted in conditions such as bipolar disorder, schizophrenia and dementia [7, 8]. In humans, twenty-four-hour rhythms in the blood transcriptome and physiological variables are also disrupted when the sleep-wake cycle and associated feeding-fasting cycles are desynchronised from the SCN such as occurs during shiftwork and jet lag, or when participants carry a sleep debt [9–11]. In both humans and rodents, this disruption can vary across different organs and tissues [12, 13] and is likely to mediate some of the negative health consequences of, for example, shift work [14]. The disruption of rhythmicity in tissues and organs may be limited to rhythmicity downstream from the core circadian molecular clock, or encompass disruption to the core molecular clock itself.

Quantifying tissue- and organ-specific rhythmicity in humans and other species will provide new insights into the temporal organisation of gene expression and holds promise for circadian medicine [15, 16]. Quantifying rhythmicity in tissues and organs requires time series of samples, but for humans time series samples of most tissues and organs cannot easily be obtained. One tissue that is easily accessible in living humans is blood. It has been shown that the human blood transcriptome contains information on altered gene expression profiles in for example the brain [17] and lung [18]. Recently, machine learning approaches demonstrated that the human blood transcriptome can be used to partially ‘predict’ the transcriptome in 16 out of 26 analysed tissues [19]. These studies used single time point samples and could not assess whether rhythmic aspects of the human blood transcriptome are concordant with rhythmicity of gene expression profiles in tissues and organs. Analyses of human blood transcriptome time series have already demonstrated that rhythmicity in sets of blood transcripts reflect the phase of the human SCN, as indexed by the melatonin rhythm [9,20–22]. However, to what extent rhythmicity in the human blood transcriptome overlaps with rhythmicity in other brain areas, tissues and organs remains unknown. Time series of tissue-specific transcriptomes for a large number of human tissues and organs are not available and comparison of the human blood transcriptome with available transcriptome time series of tissues and organs in rodents [23, 24] may be less informative and relevant, in part because rodents are nocturnal and are, from an evolutionary perspective, quite distant from humans. Time series of transcriptomes of sixty-four tissues and organs of a diurnal non-human primate, the baboon, have recently become accessible [25]. In this species, across all tissues/organs, 81.7% of protein-coding genes were rhythmic and highly tissue/organ-specific, such that no single gene was rhythmic in all tissues/organs, and a total of 3,199 baboon genes were rhythmic in only one tissue/organ. Most of the rhythmic genes were universally expressed genes, i.e. genes expressed (but not necessarily rhythmic) in all tissues/organs, supporting basic processes like transcription, RNA processing, DNA repair, protein homeostasis and cellular metabolism. Analysis of the timing of rhythmicity in the baboon tissues and organs revealed, similar to the human blood transcriptome [26, 27], a striking bimodality with peaks in the early morning and late afternoon. Analysis of the timing of core clock genes indicated that in the baboon their timing was rather similar across tissues and organs but differed from their timing in the nocturnal mouse.

Here, we compared the similarities in expression and timing of human blood and baboon tissue/organ-specific transcriptomes to determine: 1) the overlap between rhythmic transcripts in human blood and transcripts shown to be rhythmic in 64 tissues/organs of the baboon, 2) how these rhythms are aligned with light dark cycles, and 3) whether timing of rhythmic transcripts in baboon tissues and organs is similar to their timing in human blood. A flow diagram illustrating the various analyses and comparisons we have conducted can be found in Supplemental Figure 1.

## Methods

### Human and Baboon data sources

#### Human

Human blood transcriptome data collected in protocols approved by local ethics committees and as previously described [26, 27] were accessed via GEO (www.ncbi.nlm.nih.gov/geo) [28, 29] (accession no. GSE48113 and GSE39445, respectively). The first dataset represents the baseline <u>diurnal</u> condition in which 22 participants were scheduled to sleep during the night for 9 h and 20 min in phase with the circadian clock. The second dataset represents the sleeping out-of-phase with the circadian clock condition. The same 22 participants continued on a forced descinchrony protocol to achieve a sleep-wake cycle where the sleep period is 12 h out of phase with the melatonin rhythm. Seven 4-hourly blood transcriptome samples (spanning > 24 h) were collected in each condition, with sampling starting 4 hours (for sleeping in-phase) and 16 hours (for sleeping out-of-phase) before the end of the light phase [27]. For this analysis, we extracted all baseline (in-phase) and mistimed (out-of-phase) samples for 19 participants (all participants excepting those labelled 104, 141, and 219, due to a lack of samples passing QC for time series analysis, as described in [27]), and performed quantile normalisation. The third dataset represents the human <u>circadian</u> condition; data from 26 participants were collected under constant routine conditions, i.e. during approximately 40 h of wakefulness in dim light, immediately following 1 week of exposure to a 14 h light 10 h dark cycle and sufficient sleep [26]. Here, we extracted the data for the ten 3-hourly blood samples collected from 19 participants (all participants excepting those labelled 79, 91, 112, 182, 186, 214 and 267, due to a lack of samples passing QC for time series analysis, as described in [26]), and performed quantile normalisation. Rhythmic transcripts (probes) and their acrophase (i.e. the timing of the fitted maximum of the rhythm) in human blood were as reported [26, 27] and defined as having a prevalent (across participants) oscillatory component with a ∼24-h period.

#### Baboon

Rhythmic genes, and their associated attributes (e.g. acrophase), in baboon were as provided by [25]. Rhythmicity of a gene within a specific tissue/organ was assessed by applying Metacycle [30] to the single (no biological replicates), time series consisting of 12 samples taken 2 hours apart. Circadian period was allowed to vary between 20 – 26 h. In a given tissue/organ, genes with a Metacycle P-value < 0.05 were considered to be rhythmic. The processed baboon expression data were downloaded from the Gene Expression Omnibus (GEO, accession: GSE98965 [28, 29]).

### Human and Baboon orthology

Similarity between the rhythmic transcriptome in human blood and the rhythmic transcriptome of baboon tissues/organs was assessed on the set of orthologous genes. Orthology between human and baboon genes was as defined in Ensembl [31] using R [32] with package BioMart [33] (v2.40, accessed 23/1/2019). In total, there were 13,838 unique human genes (Ensembl IDs) and 13,960 unique baboon genes assessed by both Archer et al. [27] and Mure et al. [25] that we considered (Supplemental Figure 2). Of the 1,502 unique microarray probes identified as targeting rhythmic genes in the blood transcriptome of human participants on a normal sleep-wake cycle, 843 probes, targeting 759 unique human genes (Ensembl IDs), could be mapped to 786 orthologous baboon genes assessed by Mure et al. (Supplemental Figure 2). For the overlapping rhythmic genes between human blood and across baboon tissues/organs, the overlap percentage between two sets was defined as 2*(c)*100/(n+m), where c is the number of common elements, n is the size of first set and m is the size of the second set.

### Gene expression levels of human blood and baboon tissues/organs were derived using different platforms

RNA-seq has a wider dynamic range and allows for sequencing of the whole transcriptome while microarrays only profile pre-determined transcripts. However, both platforms have their own biases. The difference in the platforms used to estimate mRNA abundance did not hinder the analyses conducted because all comparisons were performed using either z-scored gene expression temporal profiles or gene expression peak times. The z-scored gene expression temporal profiles represent the deviation of expression level relative to the mean value in standard deviation units, while the acrophase value represents the time at which the maximum expression value is estimated. Both objects represent time-dependent relative changes in expression within baboon or within human, and as such, are robust across platforms. Microarray models can directly predict RNA-seq-profiled samples if the gene-expression data were z-scored before modelling and prediction[34].

### Circular cross-correlation of expression profiles

Cross-correlation is a technique that can be used to compare two time series. The procedure identifies the displacement of one time series relative to the other that provides the best match, measured by correlation, between the two time series. The basic process involves: Calculate a correlation coefficient between the two series.

Shift one series creating a lag and repeat the calculation of the correlation coefficient.

Repeat 1 and 2 for all time points.

Identify the lag with the highest correlation coefficient.

In the circular cross-correlation the data is shifted circularly. To achieve this we translated the Matlab implementation of the circular cross correlation [38] to R. A comparison of two hypothetical time series with a four-hour lag using circular cross-correlation is shown in Supplemental Figure 3.

### Alignment of human and baboon data to the light-dark cycle

During baseline diurnal conditions, the timing of the light-dark cycles to which human and baboon were exposed were different; human 18h:40min light: 9h:20min darkness; baboon 12h light:12h darkness. When the duration of the photoperiod is different across conditions, the choice of the reference external phase marker (e.g. onset of light period, onset of dark period, midpoint of light period, clock time) has an impact on the observed alignment of circadian rhythmicity across these conditions [35]. External phase markers that have been used are the midpoint of the light period [35], sunrise e.g. [24], the onset of waketime (i.e. the onset of the light period in humans) or habitual bedtime (i.e. the end of the light period in humans) [36].

To identify the appropriate phase reference point we compared the alignment of human and baboon molecular rhythms using circular cross-correlation on three reference points: onset of light, the midpoint of light, and the end of light. Human blood gene expression profiles of each participant were z-scored and aligned to either start, midpoint or end of light, and interpolated to the baboon sampling time points with the same light alignment. For each probe targeting a human/baboon orthologous gene, the mean and standard deviation across participants per interpolated time point were calculated. The mean values were used to calculate the circular cross-correlation with the z-scored baboon gene expression profiles.

Circular cross-correlation between the z-scored expression profiles referenced to either lights on, midpoint of light, or end of light were computed. The lag at which the maximum circular cross-correlation coefficient was obtained was taken to represent the level of alignment. A lag of zero indicates no difference in 2-h binned peak times, a negative lag x indicates the transcript peaks x h in human blood before the baboon tissue, and a positive lag y indicates the transcript peaks y h in the baboon tissue before the human blood.

The lags identified using the circular cross-correlation based on alignment with start, midpoint and end of light were ranked for each probe such that the alignment with the smallest lag was given the top rank, i.e. rank 1. Ranks of lags of all rhythmic probes were averaged within each baboon tissue/organ or across all tissues/organs. The resulting averages were used for the comparison of alignment strategies.

### Comparison of human and baboon acrophases

We obtained the human blood acrophases from Archer et al. 2014 and the baboon acrophases from Mure et al. 2018. All acrophases were expressed relative to the selected phase marker (i.e. aligned with start, midpoint or end of light). All calculations involving acrophases were performed using circular statistics.

Circular correlation from the R circular package [37] [v 0.4-93] was used to compare the acrophases (i.e. the timing of the maximum relative to the phase reference point) of rhythmic transcripts in human blood and rhythmic transcripts in baboon tissues and organs. Supplemental Figure 4 describes the comparison of human and baboon acrophases using circular correlation. The human blood acrophases were represented by the average value across all participants. Benjamini-Hochberg (BH) correction [38] was applied to the P-values of the circular correlations of the tissues/organs.

### Circular clustering analysis

K-means clustering [40] was used to cluster rhythmic genes based on their acrophases in human blood and baboon tissues/organs. We implemented a circular version of the clustering method by using the Cartesian coordinates (x,y) of the acrophases as input to the clustering function, i.e. x=cos(θ) and y= sin(θ), where θ is the acrophase value in radians.

### Enrichment analysis

Over-representation of Gene Ontology (GO) biological process and molecular function terms within a given list was performed using the functional enrichment analysis tool Webgestalt R accessed in August 2020 (v0.4.3) [41]. Default settings were used except that the background set was the Agilent 4×44k v2 as used in Archer et al. [27], and the false-discovery adjusted P-value threshold was set to < 0.05.

Over- and under-representation of blood immune cells within a given set of rhythmic genes was assessed using a two-sided Fisher exact test. Immune cell-markers were as defined previously [42]. All calculations were adjusted to include only orthologous human/baboon genes.

### Identifying blood-based biomarkers for timing of tissue/organ-specific rhythmicity

We conducted a proof-of-principle exercise to explore whether the phase of rhythmicity in tissues/organs could potentially be ‘predicted’ by a selected set of transcripts from one or two blood samples. To develop and validate such a multivariate whole-blood mRNA-based putative predictor of tissue/organ-specific rhythmicity, which does not require a time series of samples, we proceeded as follows. We used a single gene marker as a tissue phase marker (See Supplemental Figure 5). This gene was selected based on the circular cross-correlation analysis between human blood and the selected baboon tissue/organ per participant. The marker was required to have a high average circular cross-correlation and a small standard deviation of circular cross-correlation lag (< 2 h) across participants. The marker gene was selected from the top candidates according to these criteria and applying a tissue/organ specificity criterion, i.e. the gene marker should be rhythmic in only a low number of baboon tissues/organs. From the remaining (non-marker, non-clock gene) transcriptome features we generated a predictive model for the timing of tissue-specific rhythmicity. We performed this exercise for two tissues: paraventricular nucleus and the antrum of the stomach. In these tissues, the correlation of the acrophases of the overlapping rhythmic genes in blood and baboon tissue was significant (BH corrected P < 0.05). The tissues were also characterised by a relatively large number of rhythmic genes shared between the baboon tissue and human blood, and the overlap of these rhythmic genes between the two baboon tissues was relatively small. A final selection criterion was that the two tissues represented different physiological systems in the baboon. The model’s target was the human blood sample phase relative to the tissue phase given by the gene marker, i.e. the model predicts the phase of rhythmicity in the tissue at which a blood sample is taken (See Supplemental Figure 5). We used a Partial Least Squares Regression (PLSR) model as described previously [22]. This model has been shown to perform equally well as other machine learning approaches [e.g. 22]. Input to the model was the set of probes (transcriptome features) targeting orthologous genes rhythmic in both human blood and the selected baboon tissues/organs, excluding canonical clock genes. Samples (N = 129) were pre-processed and split into a training set (sleeping in-phase dataset, N = 61 samples, 9 participants) to develop the models, and two independent validation sets (set 1: sleeping in-phase dataset, N = 68 samples, 10 participants; set 2: sleeping out-of-phase, N=68 samples, 10 participants) to test the predictive performance of the models as described in [22]. As a control, we also tested the predictive performance of random sets of transcriptome features targeting orthologous genes, where the size of each set was the same as the PLSR-selected set.

## Results

### Overlap of the rhythmic transcriptomes of human blood and baboon tissues/organs

652 (∼5.7%) of the 11,351 orthologous genes targeted by the human microarray platform and reported by [25] to be rhythmic in at least one of the sixty-four tissues/organs of the baboon, were also rhythmic in human blood samples collected in the presence of a sleep-wake cycle (Supplemental Figure 2, Supplemental Table 1). These 652 genes represented 86% of the orthologous genes (759) that were rhythmic in human blood (Supplementary Figure 2). 438 (∼67%) of these 652 genes were considered universally expressed genes (UEGs) in the baboon (Supp. Table S4 in [25]). The 652 human genes that were rhythmic in both baboon tissues/organs and human blood also included clock associated genes (e.g. Arntl, Nr1d2, Per2, Per3, Nfil3, Npas2, Cry2, and Tef) and these will be discussed in detail below.

Functional enrichment analysis for over-represented non-redundant GO biological processes and molecular functions in this set of 652 genes identified significantly enriched (FDR < 0.05) terms not related to blood-specific processes and functions but terms related to translation (e.g. ribonucleoprotein complex biogenesis, tRNA metabolic process, translation initiation, translation elongation). The most significant GO term was ‘structural constituent of ribosome’ (enrichment ratio 4.73, P = 4.273e-12, FDR = 4.837e-9) with an overlap between human and baboon of 28 genes for ribosomal subunits, including 9 genes for mitochondrial ribosomal subunits.

### Distribution of all overlapping rhythmic genes across tissues/organs

The overlap of rhythmic genes in tissues/organs with other organs/tissues in the baboon and with rhythmic genes in human blood is illustrated in Figure 1. Across tissues/organs, an average of 6.02% ( SD 1.73%) of each individual tissue/organ rhythmic transcriptome was shared with the human blood rhythmic transcriptome and the percentage of overlap was on average 5.4 % ( SD 2.15%). This compares to an overlap of rhythmic transcriptomes of on average 8.83% ( SD 8.05%) between dyads of tissues/organs in the baboon (Fig 1 b). The percentage of genes that were rhythmic in human blood and in baboon samples were not distributed evenly across the tissues/organs (Figure 1, Supplemental Table 2). For each of the 64 tissues/organs at least one rhythmic gene was also identified as rhythmic in human blood. The gene identified as rhythmic in the greatest number of baboon tissues was Arntl (46 baboon tissues). This was followed by Nr1d2 (38 tissues), Per2 (38 tissues), Per3 (36 tissues), Nfil3 (33 tissues), Npas2 (24 tissues), Cry2 (24 tissues), and Tef (24 tissues). Although these 8 genes are all associated with the core molecular clock and were all rhythmic in human blood, they were all rhythmic in only six baboon tissues/organs (cornea, pancreas, skin, white adipose tissue, thyroid and retinal pigment epithelium).

**Figure 1:**
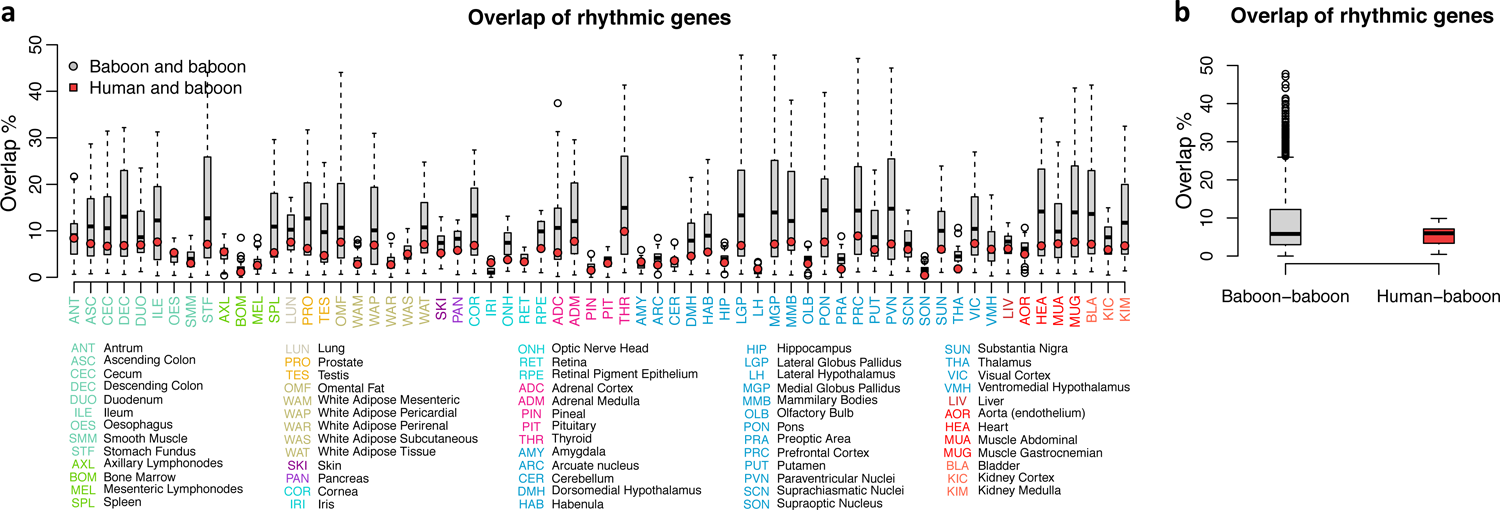
Overlap of genes that are rhythmic in baboon tissues/organs and human blood. a) Overlap of rhythmic genes between blood and specific tissues (red dots), and overlap of rhythmicity in tissues/organs and other tissues/organs (gray box plots). b) Average overlap between dyads of baboon tissues/organs and human blood with tissues/organs. Colour-coding of tissues/organs reflect the same physiological grouping as [25].

### Detecting genes that are rhythmic in only one baboon tissue/organ in the rhythmic transcriptome in human blood

For 46 of the 64 tissues/organs at least one tissue/organ-specific rhythmic gene was identified as rhythmic in human blood (see Supplemental Table 1). Of the in total 1,788 genes reported to be rhythmic in only one baboon tissue/organ, 109 (∼6%) were also identified as rhythmic in human blood collected in the presence of a sleep-wake cycle. Only 42 (∼39%) of these 109 genes were considered UEGs related to basic biological functions, e.g. translation (Rpl36a, Mrpl47, Mrps33), lipid metabolism (Abhd5, Fabp5, Acsm3), antioxidant activity (Alox5ap, Gpx7), small molecule metabolism (Clybl, Dnajc15, Pask).

Examples of the time course and alignment of tissue/organ-specific rhythmic genes are plotted in Figure 2. Examples were selected for genes that were robustly rhythmic and mostly coded for proteins with functions relevant for the tissue/organ. Of these example genes, only one, Aoc3, is classified as a UEG in the baboon. Some of these tissue/organ-specific genes are associated with functions that are relevant to those tissues/organs. Shisa7, which in the baboon was only classified as rhythmic in the mammillary bodies, is also rhythmically expressed in human blood both in the presence of a sleep-wake cycle and in the absence of a sleep-wake cycle with a phase difference of only −2 h (Figure 2a). Shisa7 is implicated in synaptic potentiation. Mobp (Figure 2c), classified as rhythmic only in the globus pallidus, where its expression is closely aligned with its expression in human blood with a phase difference of 4 h, is implicated in myelin sheath regulation. Another brain tissue-specific rhythmic gene is Tmem145, which was classified as rhythmic only in the paraventricular nucleus. It has no known function but its time course in the paraventricular nucleus is closely aligned (lag = 0 h) with the human blood profiles (Figure 2b). Tissue/organ-specific rhythmic genes were also observed in peripheral tissues and organs. Rnf207, a gene implicated in cardiac K channel regulation, was classified as rhythmic only in the baboon heart but was also rhythmic in human blood with a lag of 0 h (Figure 2d). Umod is a gene that is important for renal function and was only rhythmic in the kidney cortex and in human blood with a phase lag of 0 h (Figure 2e). Otx1 was rhythmic only in the cornea of the baboon as well as in human blood with a lag of −4 h (Figure 2g). This gene plays a role in sensory organ development. Aoc3 was specifically rhythmic in axillary lymph nodes, was also rhythmic in human blood with a lag of 2 h and its function is related to lymph node trafficking (Figure 2h). As for brain tissues, some peripheral tissue/organ-specific rhythmic genes were not associated with a known function (e.g., Tmem116 in the bladder Figure 2f). All tissue/organ-specific rhythmic genes that were also rhythmic in human blood are listed together with their functional annotation, where known, in Figure 3. It should be noted that none of the tissue/organ-specific rhythmic genes has any known direct connection with circadian function.

**Figure 2:**
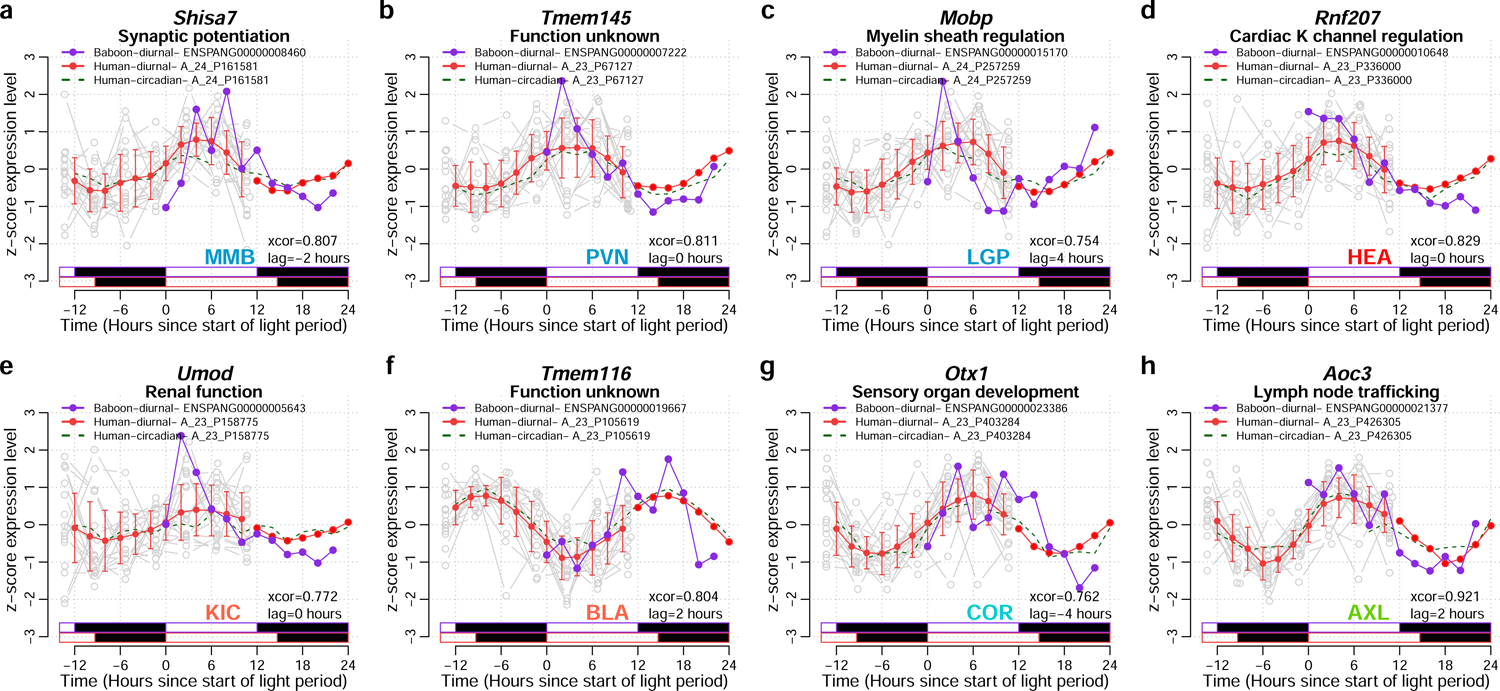
Examples of tissue/organ-specific rhythmic genes in baboon and orthologous transcripts also rhythmic in human whole blood. Tissue/organ-specific rhythmic gene examples for a) MMB, mammillary bodies, b) PVN, paraventricular nuclei, c) LGP, lateral globus pallidus, d) HEA, heart, e) KIC, kidney cortex, f) BLA, bladder, g) COR, cornea, h) AXL, axillary lymphonodes. For each tissue/organ-specific rhythmic gene (gene symbol at top of each panel), the z-scored expression profiles are plotted against time since the start of light for the baboon in purple (Ensembl ID for baboon genes provided in symbol legend) and for the orthologous human transcript in solid red line (mean +/- SD for interpolated data points and just mean for double plotted data points, actual data points for each participant are shown in gray) for the baseline diurnal plots, and as a dashed green line for the circadian plots (human transcript Agilent probe IDs provided next to symbol legends). In each panel, the strength of circular cross-correlation between the alignment of the baboon expression profile and the human diurnal expression profile is listed (xcor), together with the lag in hours between the expression profiles. The relative light/dark cycles for baboon (purple) and human diurnal (red) conditions are indicated as horizontal bars below the plots (white = light, black = dark). For each gene, the function of its coded protein (where know) is listed below the gene symbol.

**Figure 3:**
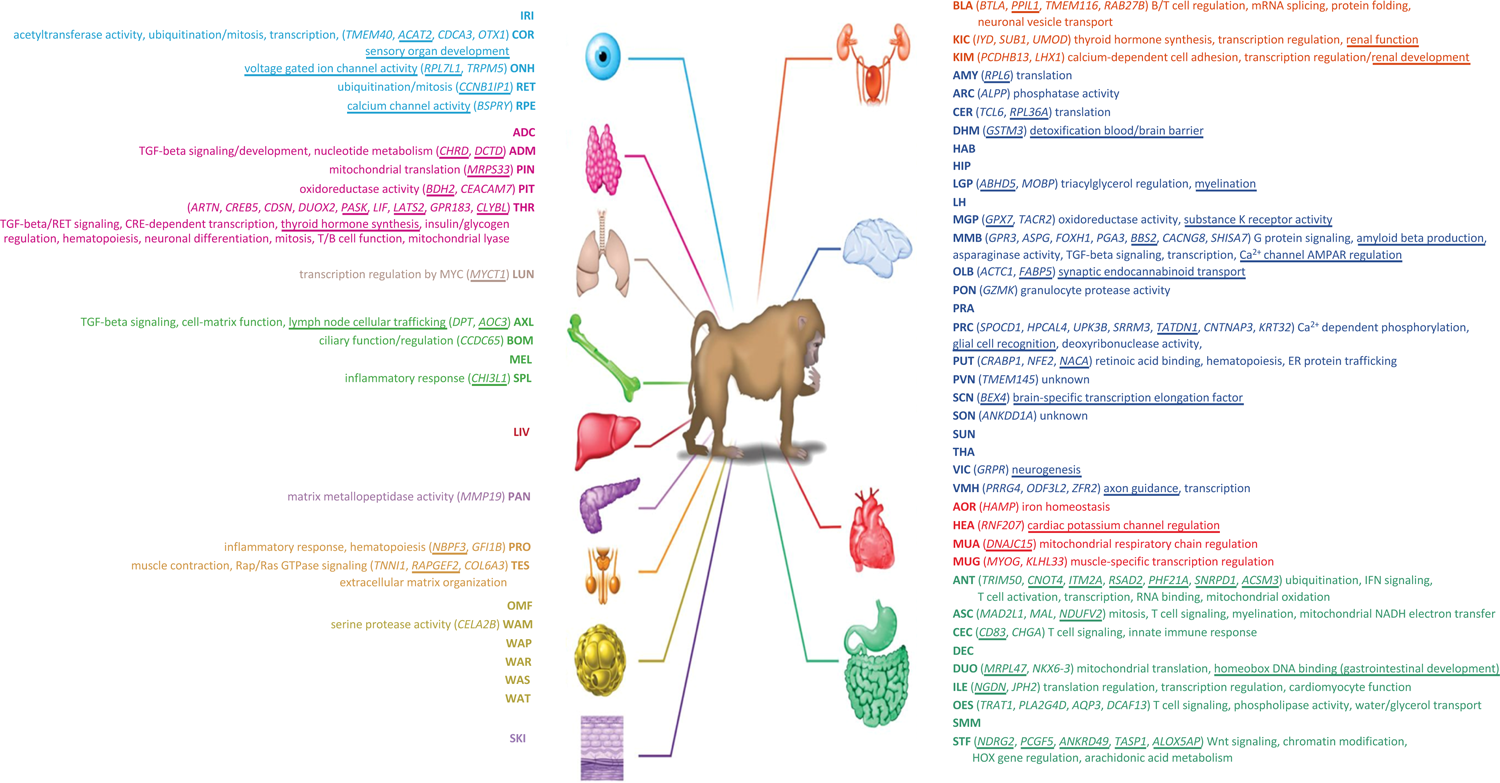
Tissue and organ-specific rhythmic genes in baboon and human blood. For each baboon tissue/organ, genes that are only rhythmic in that tissue/organ and also rhythmic in human blood are listed. Underlined genes are genes that are universally expressed in the baboon. Known protein functions associated with the genes are also listed. Underlined protein functions are directly related to the tissue/organ. Figure adapted from [25] (with permission) and the same tissue/organ colour scheme has been used. IRI iris, COR cornea, ONH optic nerve head, RET retina, RPE retinal pigment epithelium, ADC adrenal cortex, ADM adrenal medulla, PIN pineal, PIT pituitary, THR thyroid, LUN lung, AXL axillary lymphonodes, BOM bone marrow, MEL mesenteric lymphonodes, SPL spleen, LIV liver, PAN pancreas, PRO prostate, TES testis, OMF omental fat, WAM white adipose mesenteric, WAP white adipose pericardial, WAR white adipose perirenal, WAS white adipose subcutaneous, WAT white adipose tissue, SKI skin, BLA bladder, KIC kidney cortex, KIM kidney medulla, AMY amygdala, ARC arcuate nucleus, CER cerebellum, DHM dorsomedial hypothalamus, HAB habenula, MMB mammillary bodies, OLB olfactory bulb, PON pons, PRA preoptic area, PRC prefrontal cortex, PUT putamen, PVN paraventricular nucleus, SCN suprachiasmatic nuclei, SON supraoptic nucleus, SUN substantia nigra, THA thalamus, VIC visual cortex, VMH ventromedial hypothalamus, AOR aorta, HEA heart, MUA muscle abdominal, MUG muscle gastrocnemian, ANT antrum, ASC ascending colon, CEC cecum, DEC descending colon, DUO duodenum, ILE ileum, OES oesophagus, SMM smooth muscle, STF stomach fundus.

### Diurnal vs Circadian overlapping rhythmic genes

Of the 652 overlapping genes that were rhythmic in the presence of a sleep-wake cycle, only 152 genes (23.3%) were also rhythmic in the absence of a sleep-wake cycle. These truly circadian, rather than just diurnal genes, were observed in all tissues/organs apart from bone marrow and the supraoptic nucleus (Supplemental Table 1). The highest number of truly circadian human overlapping genes was found in the thyroid (76 probes targeting 60 genes). GO enrichment analysis of the 152 truly circadian genes identified non-blood-specific processes such as ribonucleoprotein complex biogenesis and cellular carbohydrate metabolic process, but unlike the larger 652 gene set, also blood-related processes such as negative regulation of cytokine production, type 2 immune, and 0.05 (see Supplemental Table 3).

### Diurnal vs Circadian for tissue-specific overlapping rhythmic genes

Of the 109 overlapping tissue/organ-specific genes that were rhythmic in the presence of a sleep-wake cycle, only 25 (22.93%) were also rhythmic in the absence of a sleep-wake cycle. These truly circadian, rather than just diurnal tissue-specific genes were associated with 14 tissues. Tissues with the highest number of circadian genes were thyroid (Clybl, Pask, Creb5, Lats2), prefrontal cortex (Hpcal4, Krt32, Cntnap3, Spocd1), oesophagus (Aqp3, Trat1), cornea (Otx1, Tmem40) and antrum (Phf21a, Rsad2) (Supplemental Table 1). Some of these tissue-specific circadian genes have functions related to the tissue, e.g. Otx1 (orthodenticle homeobox 1) expressed in the cornea regulates brain/sensory organ development, and Cntnap3 (contactin associated protein family member 3) expressed in the prefrontal cortex and mediates neuron-glia interaction.

### Alignment of human blood and baboon rhythmic genes

#### Alignment of rhythmic expression profiles to the light-dark cycle

To optimally compare the timing of rhythmic expression profiles in baboon tissues/organs to human blood, we first identified which phase reference point yielded the closest alignment between the baboon and human transcriptomes. For Per3 in the bladder, the lag between human and baboon was 0 h when lights-on (Figure 4a), −2 h for the midpoint of light (Figure 4b), and −4 h (Figure 4c) when end of light was taken as a phase reference point. When alignment of the rhythmic transcriptome in each of the 64 tissues/organs was considered, this superiority of lights-on as a phase reference point was confirmed (paired Wilcoxon rank sum test; lights on < midpoint of light lags P = 1.06e-06, lights on < end of light lags P = 2.787e-11) (Figure 4d,e). In the vast majority of tissues/organs (n=51 out of 64), start of light period produced the smallest lags. Exceptions included white adipose retroperitoneal tissue, paraventricular nucleus and the pons (Figure 4d). Similar results were obtained when the analyses were applied to individual rhythmic gene profiles of all tissues and organs combined (Supplemental Figure 6). All data and phase relations are therefore presented relative to lights-on.

**Figure 4:**
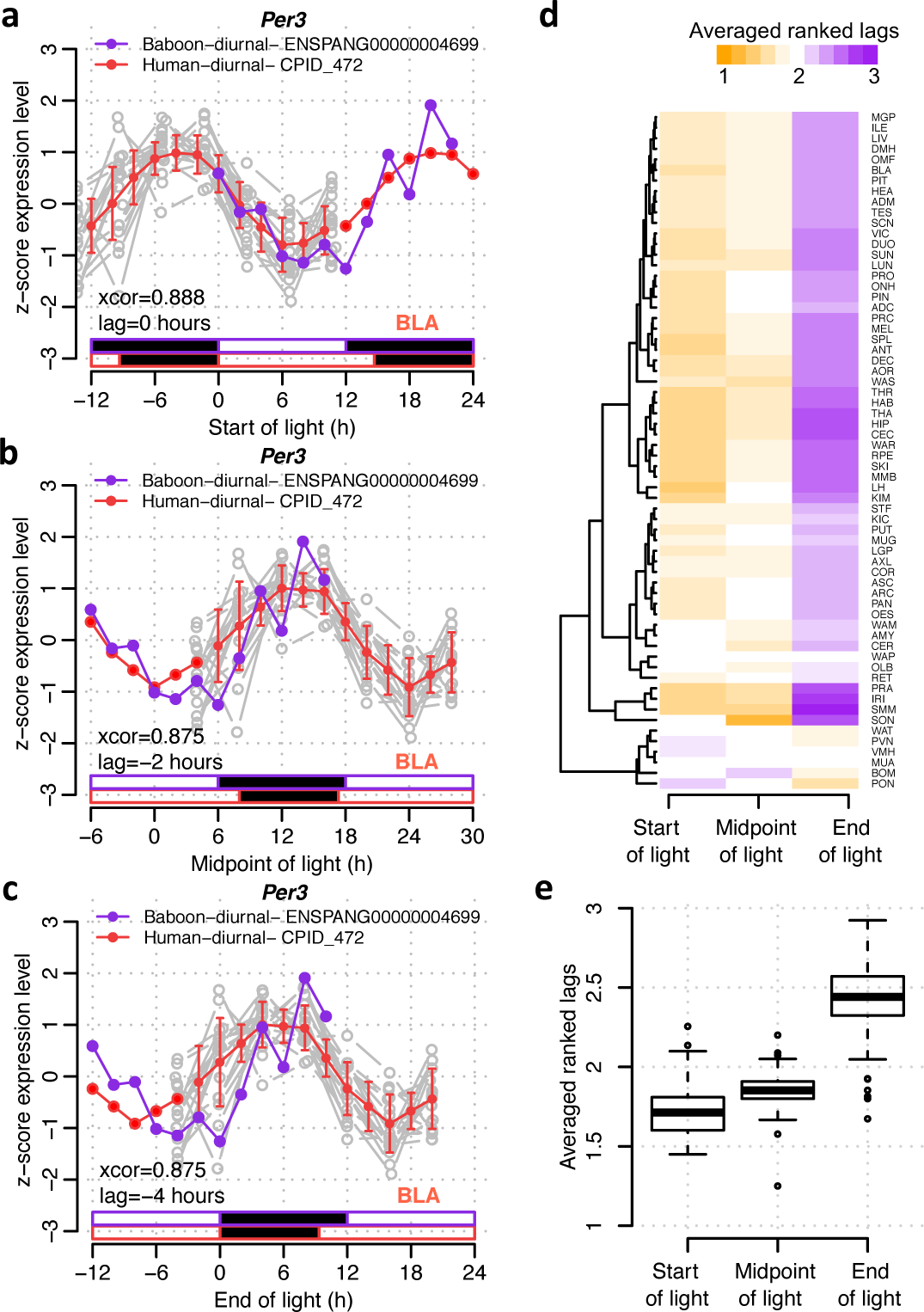
Effect of external phase reference marker on phase relationship in baboon and human blood transcriptome. For Per3 in baboon bladder and human blood, the z-scored expression profiles are plotted against time since a) the start of light, b) the midpoint of light period, and c) end of light, for the baboon in purple and for human blood in red (mean +/- SD for interpolated data points and just mean for double-plotted data points) and actual data points for each participant are shown in gray. In each panel, the strength of circular cross-correlation between the alignment of the baboon expression profile and the human diurnal expression profile is listed (xcor), together with the lag in hours between the expression profiles. The relative light/dark cycles for baboon (purple) and human diurnal (red) conditions are indicated as horizontal bars below the plots (white = light, black = dark). Lags were calculated for all overlapping rhythmic genes in every baboon tissue/organ based on alignment with start of light, midpoint of light and end of light. The resulting three lags per gene, per baboon tissue, were ranked such that the alignment with the smallest lag is given the top rank (1). d) Heatmap and e) boxplot of averaged ranked lags of all overlapping rhythmic genes, where ranking was performed within gene across the three different alignments and average was calculated for all genes within a baboon tissue/organ.

#### Alignment of ‘core clock genes under diurnal conditions’

The 652 human genes that were rhythmic in both baboon tissues/organs and human blood included clock genes. Timing of Per2, Nr1d2, Per3, Cry2 and Npas2 in human blood was close to their timing respectively, and 4.81 h, 4.63 h, 0.15 h earlier for Nr1d2, Per3 and Cry2, respectively. Nfil3 and Tef peaked on average 6.66 h, and 8.182 h, respectively, earlier in human blood than in baboon tissues/organs. Timing of Arntl profiles was on average 10 h phase advanced in human blood compared to baboon tissues/organs (lag range across tissues: −10 to +12 h) (Figure 5, and Supplemental Table 4). For Per2, Nr1d2, Per3, Cry2 the phase relationships between baboon tissues/organs and human blood were not systematically different when rhythms in the baboon brain were compared to rhythms outside the brain. Since these 8 clock genes are rhythmic in several tissues, their timing in human blood can only provide limited information about differences in timing of rhythmicity across individual tissues and organs.

**Figure 5:**
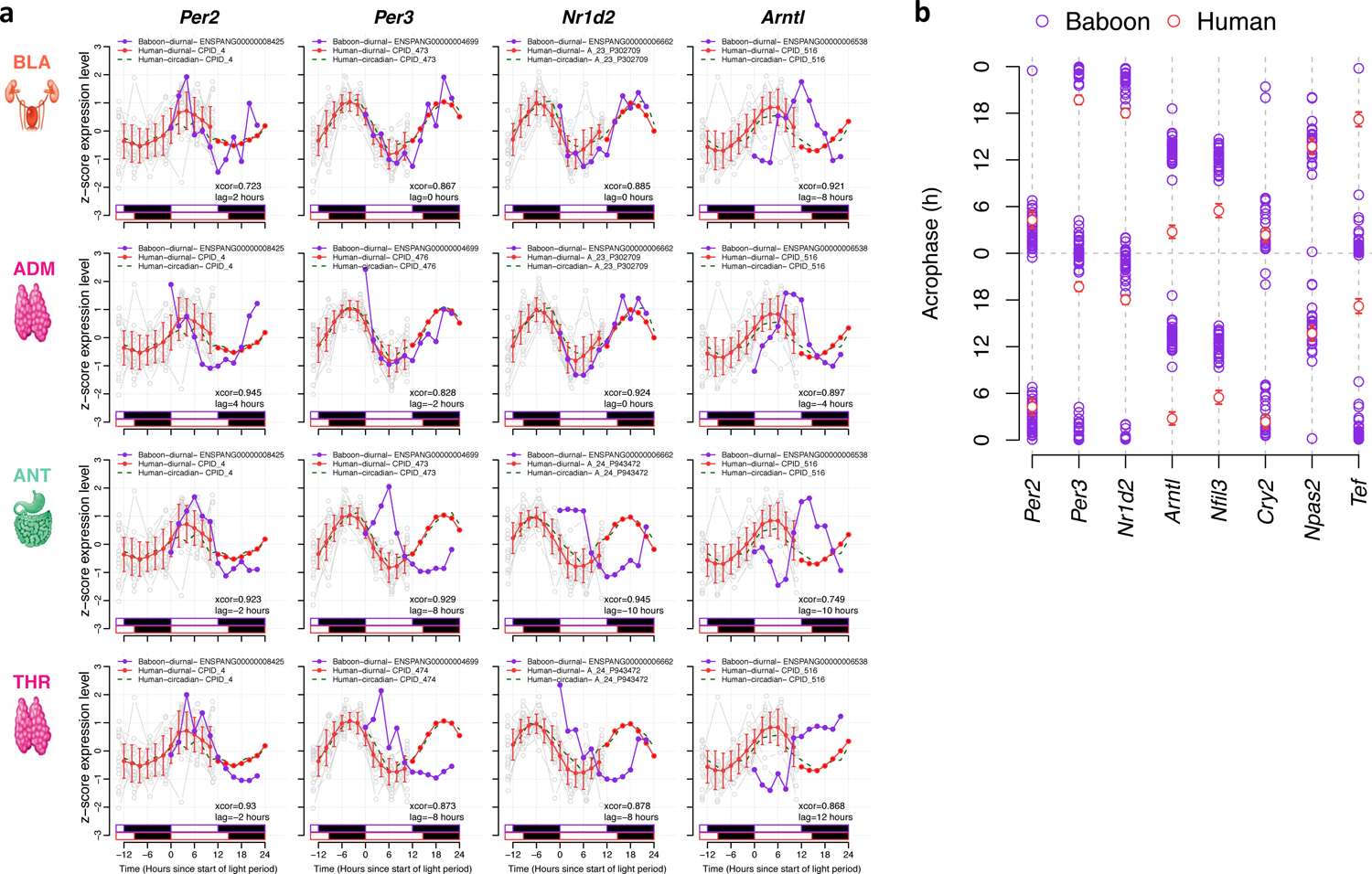
Timing of core clock genes across baboon tissues and human blood. a) Time course of example core clock genes in human blood and selected organs of the baboon. The z-scored expression profiles are plotted against time since the start of light for the baboon in purple and for the orthologous human transcript in red (mean +/- SD for interpolated data points, just mean for double-plotted data points, and gray for actual data points for each participant) for the baseline diurnal plots, and as a dashed green line for the circadian plots. b) Acrophase of clock genes in human blood and baboon tissues/organ. Acrophase values in human blood are represented by the circular mean across participants and error bars show one standard error of the mean. Please note that the data are double-plotted and acrophase values are shown relative to the start of the light period.

#### Timing of rhythmic genes in the baboon and human blood

When the timing of all overlapping rhythmic genes was considered a clear bimodality in the timing of acrophases was observed. For example, in the thyroid bimodality was observed in both human blood and the baboon when all overlapping rhythmic genes were considered (Figure 6 a, b). In human blood, the two peaks were located 2 – 4 h after lights on and 2 – 4 h after the onset of darkness. In the baboon, these two peaks were located at 6 – 10 h after lights on and in the middle of the dark phase (18 – 22 h). When all individual acrophases of rhythmic genes in baboon thyroid were plotted against the corresponding acrophases in human blood a significant association emerged (Figure 6c). Similar patterns can be observed in the mammillary bodies (Figure 6d,e,f) and antrum (Figure 6g,h,i). To further assess the extent the human blood transcriptome mirrors the baboon tissue/organ transcriptome, we used circular K-means clustering analysis to characterise the observed bi-modal acrophase distribution of rhythmic genes. We clustered all rhythmic genes into two groups based on both, their acrophase in human blood and their acrophase in baboon tissue/organs (see Figure 7a). This allowed the classification of the rhythmic genes within the two peaks of the acrophase bi-modal distributions (see Figure 7 and Supplemental Table 5).

**Figure 6:**
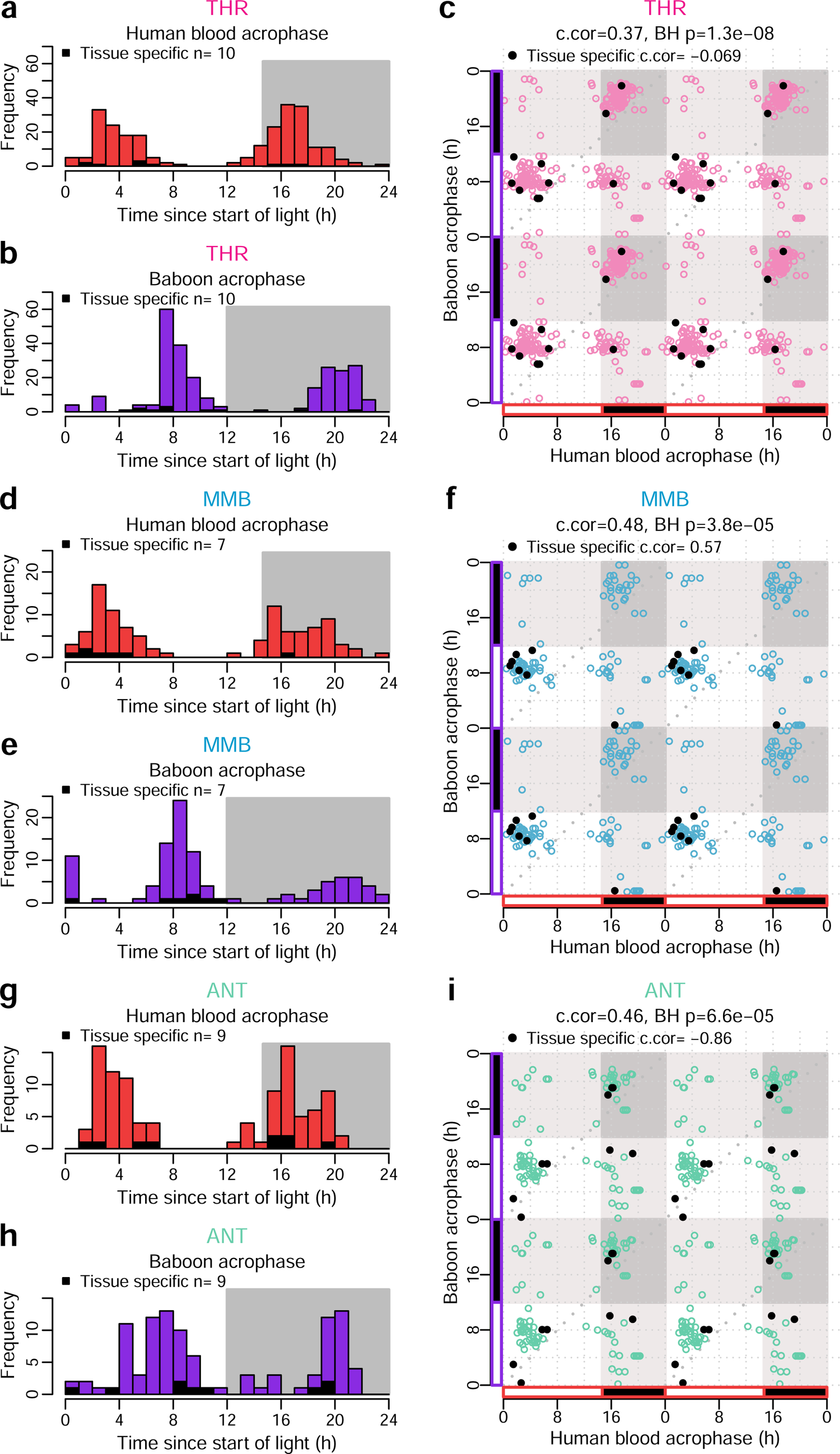
Timing of acrophases of overlapping rhythmic genes in baboon tissues/organs and human blood. Left panels: Human blood (top) and baboon (bottom) acrophase distribution of all genes that were rhythmic in human blood and in the baboon tissue/organ shown (THR, thyroid; MMB, mammilary bodies; ANT, antrum). Right panels: Scatter plot of acrophases of human and baboon genes. Open circles, all overlapping genes; filled circles, tissue/organ-specific rhythmic genes. Data are double-plotted. c-cor = circular correlation of acrophases (baboon vs. human). BH p = Benjamini-Hochberg corrected p value.

**Figure 7:**
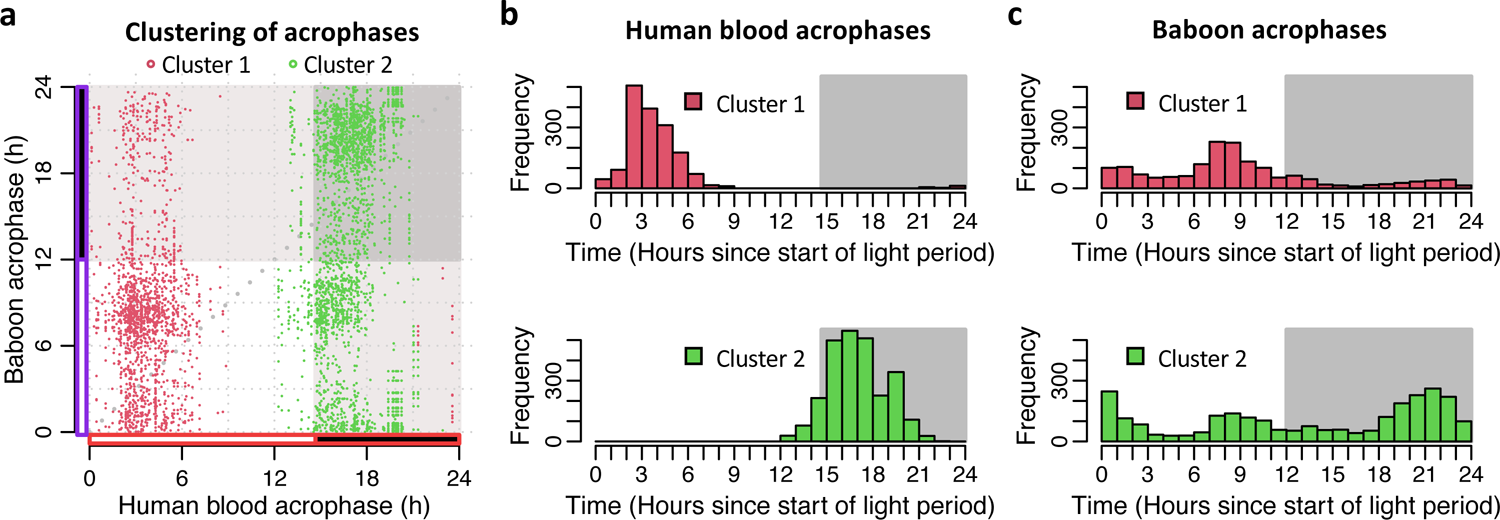
Circular clustering analysis was used to characterise the acrophase distribution of rhythmic genes in human blood and baboon tissues/organs. Clustering of all overlappingrhythmic genes into two groups based on both their acrophase in human blood and their acrophase in baboon tissue/organs. In all panels acrophases are coloured by cluster membership and are expressed as hours since lights on. a) scatter plot of acrophases in human blood vs acrophases in baboon tissues/organs. b) distribution of acrophases in human blood. c) distribution of acrophases in baboon tissues/organs.

The two cluster analysis defined genes with peak acrophases that corresponded to the human morning peak (2 - 4 h after lights on), the human early night peak (16 - 18 h after lights on), the baboon afternoon peak (6 - 10 h after lights on), and the baboon late night peak (18 - 22 h after lights on). Bimodality was also observed for the tissue/organ-specific rhythmic genes (see Supplemental Figure 7).

Genes in the night cluster for each species were strongly enriched for biological processes and molecular functions related to translation and protein synthesis (see Table 1).). Although the number of unique genes analysed in the night and day clusters were similar (286 night vs. 275 day), the greater range of processes and functions associated with day peaking genes led to reduced overall enrichment compared with the night peak. In fact, enrichment of genes for the day peak for each species did not survive the FDR < 0.05 threshold. When ranked by nominal statistical significance, these gene sets appeared enriched for a wider range of processes and functions and generalised cellular activity (e.g., extracellular structure organisation, P = 0.0002; glutamate receptor signalling pathway, P = 0.0005; organ growth, P = 0.0006; regulation of cell morphogenesis, P = 0.0029). Of interest, the day cluster contains the core clock genes Arntl, Per2, Cry2, Csnk1e, Rara, Nfil3, while the night cluster contains Per3, Nr1d2, Npas2, Tef. The night cluster also contained Rbm3, which is a temperature sensitive RNA binding protein known to regulate the translation of circadian proteins [43]. It is also of interest that genes in the human blood day cluster were not associated with typical blood specific functions such as inflammatory responses and immune function [26].

**Table 1:** Top ten significant gene ontology biological process and molecular function gene sets enriched within the night peaking baboon and human rhythmic genes.

| Gene Set | Description | P value | FDR |
| --- | --- | --- | --- |
| GO:0022613 | Ribonucleoprotein complex biogenesis | 0 | 0 |
| GO:0003735 | Structural constituent of ribosome | 0 | 0 |
| GO:0034470 | ncRNA processing | 7.7716e-16 | 2.9325e-13 |
| GO:0016072 | rRNA metabolic process | 2.6913e-11 | 7.6165e-9 |
| GO:0070972 | Protein localization to endoplasmic reticulum | 1.4028e-9 | 3.1759e-7 |
| GO:0006399 | tRNA metabolic process | 1.1740e-8 | 0.0000022150 |
| GO:0006413 | Translation initiation | 2.1438e-8 | 0.0000034669 |
| GO:0006414 | Translation elongation | 5.2676e-8 | 0.0000074537 |
| GO:0006605 | Protein targeting | 1.7375e-7 | 0.000021854 |
| GO:0140053 | Mitochondrial gene expression | 4.7592e-7 | 0.000053874 |

We next quantified the extent to which rhythmic profiles of genes in baboon tissues/organs were synchronised with the orthologous rhythmic profile in human blood by circular correlations of acrophases for overlapping rhythmic transcripts, separately for each tissue/organ. A significant correlation (BH corrected p value <0.05) was observed in 25 out of 64 tissues/organs (See Figure 8a and 8b). These 25 tissues/organs included endocrine organs such as the thyroid, brain structures such as the mammilary bodies, paraventricular nucleus and medial globus pallidus, as well as the intestine (cecum and duodenum) and the immune system (axillary lymph nodes).

**Figure 8:**
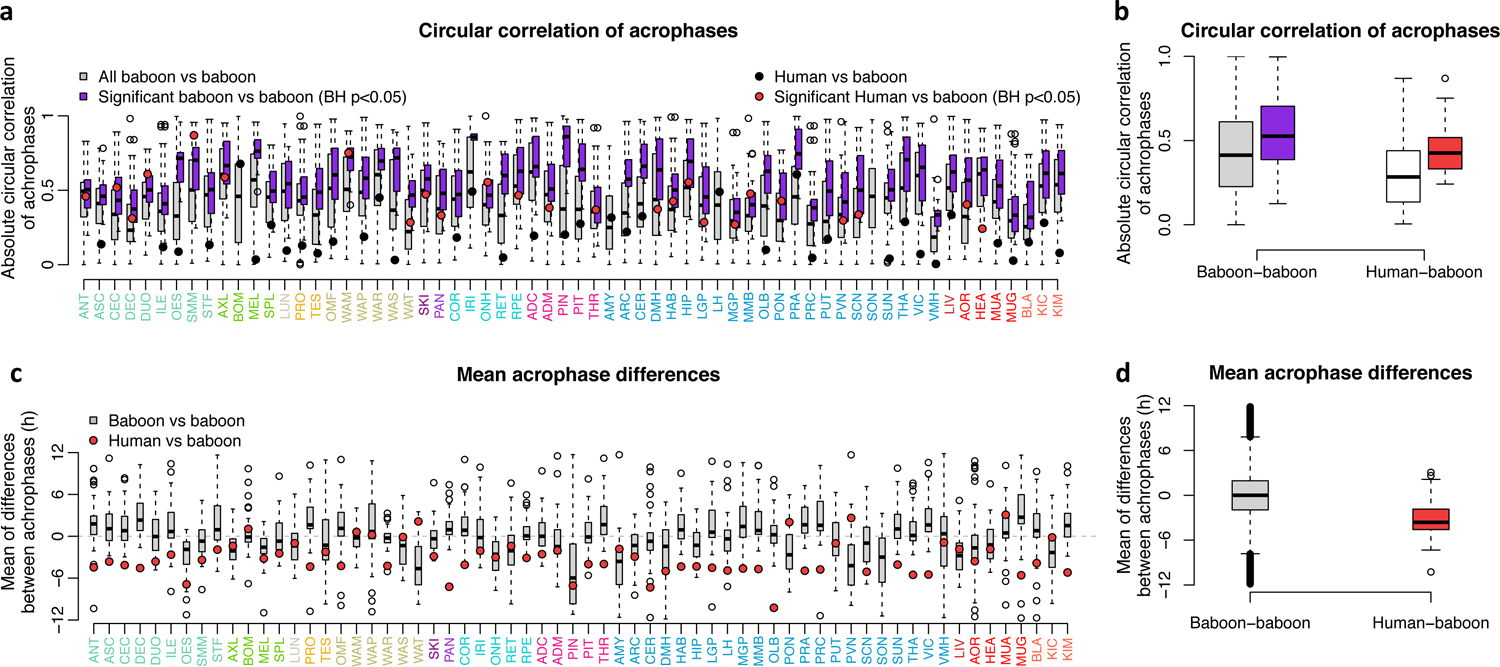
Acrophase comparison of rhythmic genes in baboon tissues/organs and their human rhythmic orthologs in blood. a) Circular correlation of acrophases of overlapping rhythmic genes for each of the 64 tissues/organs. Circles represent correlation between acrophases of the sets that overlap in blood and baboon tissues/organs; box plots represent the correlation between the overlapping sets of the tissue/organ and other tissues/organs of the baboon. b) Average circular correlation of acrophases. c) Mean acrophase differences. d) Box plot for mean acrophase differences.

When the differences in acrophases of the overlapping rhythmic genes in human blood and baboon tissues were computed, it emerged that across the 64 tissues/organs vs the human blood transcriptome human genes were timed 3.21 hours (SD 2.47 h) earlier than baboon genes. When a similar comparison was made for the timing of the overlapping transcriptomes for all baboon tissue/organ dyads the average acrophase difference was 0.00 hours (SD 3.43 h) (see Figure 8c and 8d). Similar results were obtained when we computed the circular cross correlation of the temporal profiles in baboon tissues/organs and human blood (see Supplemental Figure 8). This correlation was substantial (mean > 0.7) for most tissues/organs, with the exception of the supraoptic nucleus (Supplemental Figure 8b). Based on the circular cross correlation of the profiles, human rhythms were advanced on average by 2.69 hours (SD 2.21 h), which is less than the 3.21 hours (SD 2.47 h) estimated based on acrophase values, but it should be noted that the standard deviation of the lag was considerable for most tissues/organs (Supplemental Figure 8d).

### A blood-based multivariate single blood sample predictor for tissue specific rhythmicity?

The similarity in the timing of tissue/organ-specific rhythmicity in tissues and organs in the baboon and rhythmicity in human blood implies that the blood transcriptome may be used to assess the circadian alignment of tissues/organs. This has potential implications for the development of blood based biomarkers for rhythmicity in tissues and organs. These biomarkers may either be univariate and would require a time series of blood samples, or multivariate, i.e. based on a set of transcriptomic features which potentially would require only one or two samples.

We explored whether it was in principle possible to create a multivariate biomarker which would allow to predict the phase of tissue/organ rhythmicity from one or two blood samples. To provide proof of principle evidence, we selected the antrum of the stomach (ANT) and the paraventricular nucleus (PVN) (see Methods for selection procedure). The time of tissue-specific rhythmicity assigned to each blood sample was based on one marker gene per tissue (see Supplemental Figure 5). For the antrum we selected Slc44a2, which is a choline transporter active in the intestine, and for the PVN we selected Lrfn3, which is a synaptic adhesion-like molecule. The assumption of our analysis is that the phase of the selected blood marker gene is a good proxy for the phase of these genes in the specific tissue.

The overlap of the rhythmic transcriptome in these two tissues with the rhythmic human blood transcriptome was 81 genes and 118 genes for ANT and PVN respectively, of which only 10 genes overlapped.

To develop predictive models, we eliminated all features (targeting probes) associated with classical clock genes, because their timing is non-tissue-specific. We also excluded the selected marker gene for each tissue. The number of features selected by the predictive model were 74 genes and 113 genes for ANT and PVN respectively, of which only 8 genes overlapped (see Supplemental Table 6). From these sets of the blood-ANT and blood-PVT transcripts, PLSR applied to the training samples, identified 45 probes (targeting 41 genes) and 55 probes (targeting 49 genes) for the ANT and PVN biomarker, respectively. When these predictors were applied to the blood samples in the validation set they predicted the ANT and PVN phase with an R^2^ of 0.63 and 0.67, and a mean absolute error of 2.94 and 2.81 h for ANT and PVN, respectively (Figure 9 and Table 2).

**Figure 9:**
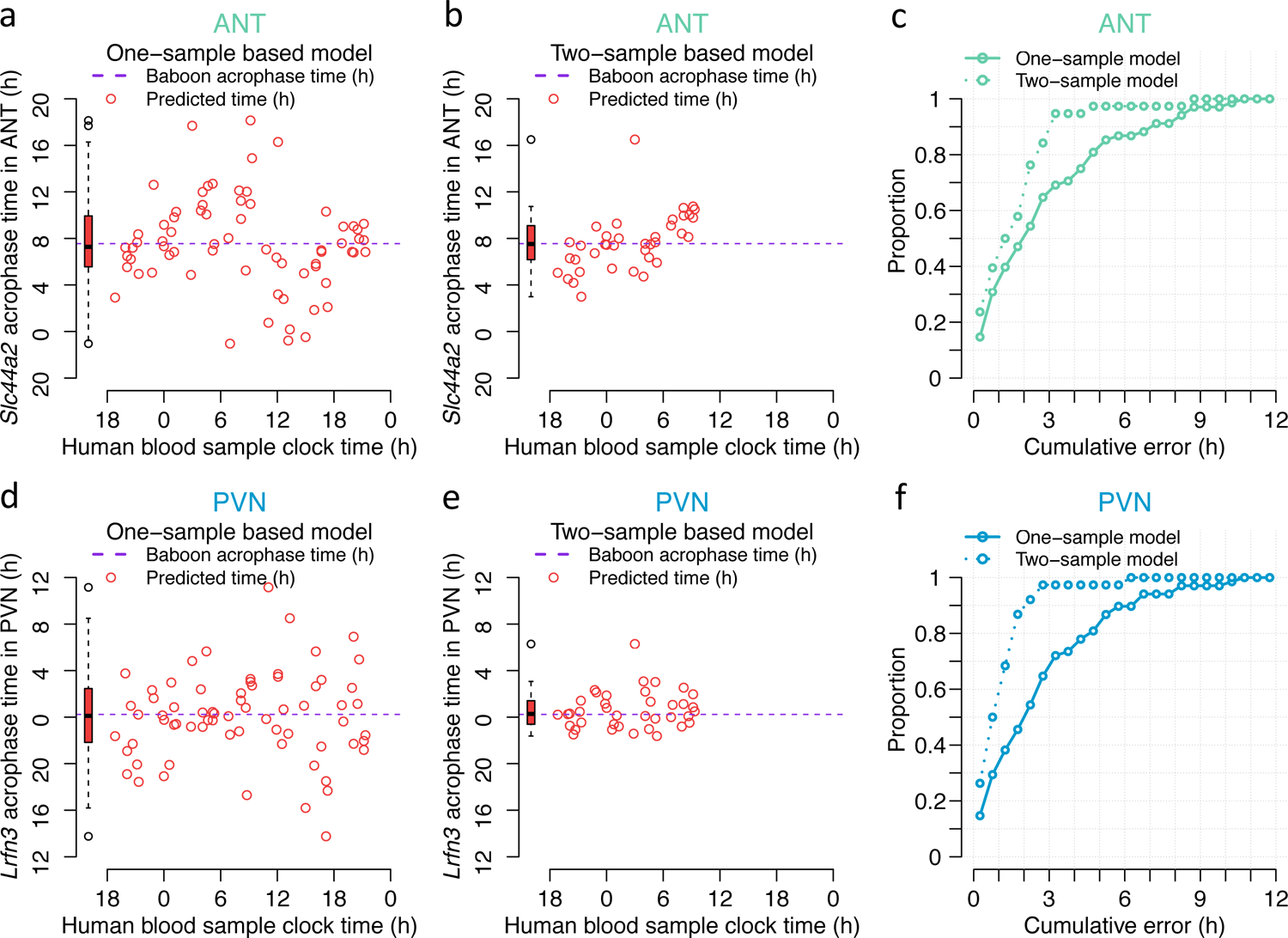
Blood-based biomarkers for tissue/organ-specific rhythmicity. Performance of one-and two-sample models derived from the training set when used to predict timing of ANT (antrum) and PVN (paraventricular nuclei) rhythmicity in the validation set. Predicted timing of ANT vs. sample clock time for each sample in the validation set for a) the one-sample and b) two-sample PLSR model. c) Proportion of predictions of timing of ANT vs. cumulative absolute error for the one- and two-sample PLSR model. Predicted timing of PVN vs. sample clock time for each sample in the validation set for d) the one-sample and e) two-sample PLSR model. f) Proportion of predictions of cumulative frequency distribution of absolute error can be used to identify the proportion of samples that can be predicted within a given error range, where the highest proportion combined with smallest error range denotes the better prediction model.

**Table 2:** Performance of phase of rhythmicity prediction in the paraventricular nucleus (PVN) and the antrum of the stomach (ANT) based on a selected set of transcripts (number of predictor genes) from one (1-sample based) or two (2-sample based) blood samples using a Partial Least Squares Regression (PLSR) model. Performance is evaluated using R2, median absolute error, error mean and standard deviation (sd), and proportion of samples with error within 1, 2 or 3 hours. The PLSR models were trained using a training set from the sleeping in-phase dataset and performance was evaluated using validation sets from the sleeping in-phase or sleeping out-of-phase datasets.

| Tissue |  | PLSR model |  | Training set | Validation set |  | Number of predictor genes | R<sup>2</sup> | Median absolute error (h) | Error mean, sd (h) | with less than |  |  |
| --- | --- | --- | --- | --- | --- | --- | --- | --- | --- | --- | --- | --- | --- |
| ANT | PVN | 1-sample based | 2-sample based | Sleeping in phase | Sleeping in phase | Sleeping out of phase |  |  |  |  | 1 h error | 2 h error | 3 h error |
| x |  | x |  | x | x |  | 45 | 0.63 | 2.19 | -0.05, 4.00 | 0.31 | 0.47 | 0.65 |
|  | x | x |  | x | x |  | 55 | 0.67 | 2.28 | 0.19, 3.77 | 0.29 | 0.46 | 0.65 |
| x |  |  | x | x | x |  | 20 | 0.71 | 1.51 | -0.13, 2.43 | 0.39 | 0.58 | 0.84 |
|  | x |  | x | x | x |  | 25 | 0.87 | 1.00 | -0.38, 1.60 | 0.50 | 0.87 | 0.97 |
| x |  | x |  | x |  | x | 45 | -0.06 | 5.60 | 0.83, 6.71 | 0.12 | 0.21 | 0.25 |
|  | x | x |  | x |  | x | 55 | -0.05 | 5.06 | 2.21, 6.36 | 0.06 | 0.18 | 0.26 |
| x |  |  | x | x |  | x | 20 | -1.20 | 6.24 | 3.58, 5.67 | 0.11 | 0.16 | 0.21 |
|  | x |  | x | x |  | x | 25 | -0.78 | 5.11 | 3.64, 4.80 | 0.11 | 0.18 | 0.29 |

For both tissues, the predictors were related to cell cycle/cell growth/apoptosis, ubiquitination/protein degradation/autophagy, and transcription/translation. Only one gene in the ANT was related to circadian rhythmicity which was *Rai1*, a transcription factor that interacts with the circadian transcription factor CLOCK.

We generated 50 negative control biomarkers for ANT and PVN by using N randomly selected probes from the set of orthologous genes assessed in Archer et al. 2014 and Mure et al. 2018, where N is the number of candidate biomarker genes (input to PLSR model) in the real model. The negative control biomarkers predicted the ANT and PVN phase with a significantly lower R^2^ (p=02).

We also investigated the performance of a two-sample based model. From the sets of the blood-ANT and blood-PVT transcripts in the training samples, the two-sample PLSR identified 20 probes (targeting 18 genes) and 25 probes (targeting 22 genes) for the ANT and PVN biomarkers, respectively. When these predictors were applied to the blood samples in the validation set they predicted the ANT and PVN phase with an R^2^ of 0.71 and 0.87, and a mean absolute error of 1.74 and 1.19 h for ANT and PVN, respectively^2^

## Discussion

### Summary of Results

Rhythmic genes in human blood overlap with rhythmic genes in baboon tissues/organs and the timing of most rhythmic genes is similar across tissues/organs and human blood.

### Extent and characteristics of overlapping rhythmicity in human blood and baboon tissues/organs

#### The sets of orthologous genes

The analyses were necessarily restricted to orthologous genes that were present in both data sets. Overall, these sets corresponded to 47% of all baboon genes as defined by Ensembl on 30/7/2020) and only 23% of all human genes as defined by Ensembl on 30/7/2020). However, here we were interested in genes that were rhythmic in both baboon and humans and this set amounted to 46.7% (652/1396) of all human blood rhythmic genes and 86% of the orthologous genes (759) in [27]. Thus, even though the overall set of orthologous genes represents only a small portion of all human genes, the set of orthologous genes that are rhythmic represents a substantial portion of the total number of human blood rhythmic genes. Improved annotation of the baboon and human genomes may allow for a more extensive comparison of the rhythmic transcriptome across human and baboon.

#### Extent and characteristics of overlapping rhythmic genes

The overlap between human blood rhythmic genes and genes that are rhythmic in one or more of the 64 tissues/organs was somewhat lower than the overlap in rhythmic gene sets across all dyads of baboon tissues/organs, but there was quite a range of overlap both across baboon tissues/organs and human blood and across baboon tissues/organs. Thus, from this perspective human blood-tissue/organ ‘dyads’ behaved similarly to any of the within baboon dyads.

The majority of rhythmic genes in the baboon and human-baboon overlapping rhythmic genes were considered UEGs in the baboon (see Supplemental Table 1). The set of overlapping rhythmic genes was primarily enriched for basic processes predominantly associated with mRNA translation, and not for specific blood functions (e.g. immune function). This indicates that the overlapping rhythmic transcriptome across blood and baboon tissue/organs is representative of rhythmicity in organs and tissues. The observation that only 39% of the set of genes that are rhythmic in human blood and in only one tissue/organ belong to the list of UEGs, and the functional annotation of this set, imply that tissue-specific rhythmic genes serve tissue-specific functions.

#### Origin of overlapping rhythmic transcripts in human blood

Human blood contains many cell types and diurnal and circadian rhythmicity in blood has been described in the number of the various cell types [46] and each of these cell types may contribute to rhythmic transcriptome in whole blood. The baboon tissues/organs were perfused prior to sampling and so were not likely to contain significant numbers of blood cells. Some tissues/organs may have contained resident immune cells, such as Kupffer cells, that cannot be removed by perfusion, but their overall contribution to the rhythmic tissue/organ transcriptome will, in general, be minimal. Therefore, the lack of, or underrepresentation of immunity-related GO terms in the human/baboon rhythmic transcript overlap is presumably due to the overall absence of blood cells in the baboon tissues/organs. We performed over/under-representation analysis of known blood immune-cell type gene markers of the overlapping rhythmic genes (see Supplemental Text S1). We found significantly higher than expected enrichment for neutrophil cell markers in some tissues, including thyroid, mammilary bodies, ileum, antrum, and bladder. Furthermore, this neutrophil enrichment was only present in the day-peaking transcripts in these tissues. These observations point to a potential albeit small contribution of this cell type to the overlapping rhythmic transcriptome in those tissues/organs. Neutrophils are the most common type of white blood cell (∼60%) and the first line of defence in the innate immune system. They are produced in large numbers during the day and then cleared by multiple tissue sites at night. In our data the neutrophil enrichment was only present in the day-peaking transcripts in these tissues and this is consistent with the rhythmicity in their abundance level which peaks during the day [46]. Some of the baboon tissue/blood overlapping gene sets that showed enrichment for neutrophil cell markers are involved with innate immune function. For example, thyroid hormones increase neutrophil function, the ileum contains Peyer’s patches which are lymphoid tissues that perform immune surveillance and signal an immune response to infection, and the antrum produces lymphoid follicles during an immune response to infection.

### Timing of rhythmicity of rhythmic transcripts in human blood and baboon tissue/organs

#### Alignment with the light-dark cycle

Baboon and humans are both diurnal species but baboons and humans living in industrialised societies are exposed to different light-dark cycles. Here, we present a first comparison of the timing of transcriptomes in these two diurnal primates. Our analyses of the alignment across tissues/organs and human blood indicates that the start of the light period, which is the activity and feeding period, is the best phase reference point for most rhythmic genes and tissues/organs, at least in the comparison of human blood and baboon tissues/organs Previous studies in which comparisons across species or conditions were made have used other phase reference points (e.g. sunrise [24]) and this may account for some of the discrepancies across these studies. Overall, these data highlight the need for the use of appropriate phase reference points for a comparison of the temporal organisation of gene expression across species and conditions.

#### Alignment of ‘clock genes’

As previously reported [22, 26], most of the known core clock genes were rhythmically expressed in human blood. In the comparison with the baboon, the most noticeable exception is Chrono (Ciart, Circadian Associated Repressor of Transcription) which was reported to be the most rhythmic gene by [25], but its human homologue, C1orf51 (targeted by two probes in the Agilent arrays), was lowly expressed and not rhythmic in human blood. This may reflect a specific functional difference in the human blood circadian clock, or it may be that the two targeting probes do not perform well. Further molecular analyses could address this.

The timing of the majority of core clock genes was similar in human blood and baboon tissues and organs. Exceptions were Arntl, Nfil3 and Tef, which were all phase advanced in human blood compare to baboon tissues/organs. The different timing of ARNTL in human blood has been noted before [24]. We note that there were 36 probes targeting ARNTL but only one of these probes was rhythmic, which may indicate that the estimate of the acrophase for ARNTL is not very reliable.

#### Alignment of all overlapping rhythmic transcripts

There are two aspects of timing that can be considered: distribution of peak times across the 24-h cycle, and the correlation of acrophases. Distribution of peak times of rhythmic genes has been reported to be bimodal in human blood [26], skin [49], mouse liver [50] and baboon tissues [25] including rhythmic splice isoforms of the same gene [51]. The exception to this is the distribution of peak times in human organs as reconstructed from samples that were not labelled with clock time [52]. In the current study, this bimodality was similarly observed in the subset of rhythmic baboon genes that were also expressed rhythmically in human blood. This indicates that the timing of these rhythmic genes in human blood is similar to the timing in baboon tissues/organs. Furthermore, for 25 out of 64 tissues/organs, the timing of the baboon genes, i.e. their acrophases, correlated significantly. Recently, several reports have discussed the associations of gene expression across tissues [25,44,45] and machine learning approaches have demonstrated correlations between gene expression levels in blood and a range of other tissues/organs [19]. Tissue-specific gene expression is not driven by a set of tissue-specific transcription factors but rather combinations of non-tissue-specific transcription factors functioning in tissue-specific regulatory pathways [44]. Thus, it is possible that blood cells share regulatory pathways that are specific to other tissues/organs.

### Diurnal vs Circadian

Many of the overlapping rhythmic genes in human blood assessed under diurnal conditions were no longer rhythmic under circadian conditions. This implies that the rhythmicity of this set of transcripts is driven by external and behavioural rhythmicity rather than circadian mechanisms which persist under constant conditions. This is consistent with many recent findings in both humans and rodents (see [53, 54]). For most practical purposes, e.g. describing abnormal rhythmicity, the diurnal condition will be most relevant. When we are interested in mechanisms underlying abnormal rhythmicity, the comparison diurnal vs constant conditions becomes relevant.

### Implications for development of blood transcriptome based biomarkers for organ and tissue specific rhythmicity

Our proof of principle exercise demonstrates that sets of rhythmic genes (associated transcriptome features) in human blood can predict the phase of marker genes in the brain (PVN) and body (Antrum of the stomach) with reasonable accuracy (mean absolute error less than 3 h, based on one blood sample) which is comparable to the accuracy of predictors of, for example, rhythmicity in skin from a skin biopsy [49].

Functional annotation of the genes that predict the phase of the PVN and Antrum (see Supplemental Text S2) did not suggest an obvious mechanism by which blood could be synchronised with the target tissue/organ. This is in contrast to the functional annotation of blood transcriptome predictors of SCN phase, which identified glucocorticoid-dependent regulation of expression as the likely mechanism of synchronisation. Blood perfuses all tissues/organs and so cross-talk between blood and tissues/organs is inevitable. Thus, it is to be expected that other blood-borne signalling agents derived from either blood or tissues/organs could synchronise gene expression in both blood and tissues/organs. [55]. Thus, blood and tissues/organs throughout the body are intrinsically linked in biological communication by a variety of mechanisms [56–58] and it is not surprising that some features of their respective transcriptomes like expression levels [19] or timing (current data) are correlated and can be used as predictors.

To further test the validity and robustness of these blood based predictors will require the availability of data on the rhythmic transcriptome of organs and tissues under conditions or in populations in which these rhythms are disrupted [16].

Our analyses already indicate that the overlapping rhythmic transcripts are affected by the presence or absence of sleep-wake and light-dark cycles, i.e. the differences between the diurnal and circadian transcripts. None of the 45 probes (targeting 41 genes) that were part of the predictor set of the ANT that were rhythmic when sleeping in phase remained rhythmic when sleeping out of phase. Only two probes (targeting SLC22A4 and MPZL1) out the 55 probes (targeting 49 genes) that were part of the predictor set of the PVN that were rhythmic when sleeping in phase remained rhythmic when sleeping out of phase. Not surprisingly, when the predictor sets were used to ‘predict’ ANT and PVN phase by using blood samples collected during sleep out of phase their ‘performance’ deteriorated drastically compared to the sleeping in phase condition (see Supplemental Figure 9 and Table 2). Whether this deterioration reflects reduction in true performance of the predictor or reduced rhythmicity in the PVN and ANT is obviously unknown because no data are available on rhythmicity in the PVN and ANT in the baboon sleeping out of phase.

### Limitations

Although the presented analyses are based on the best currently available data sets, these data sets have their limitations. The baboon data set contained no biological replicates and consisted of only 12 samples with a 2-h resolution. It may be argued that for the detection of rhythmicity these limitations lead to false positives and false negatives although the high temporal resolution (2 h) is beneficial for the accurate estimate of phase. We conducted an analysis of how error prone the cycling detection in the baboon data is likely to be and found that the data set is sufficient to accurately estimate rhythmicity and hence our estimates of the overlap of baboon and humans rhythmicity are likely to be accurate (see Supplemental Text S3).

The human data were obtained in healthy young participants. Thus, the extent to which the current results can be extrapolated to conditions in which rhythmicity in organs and tissues is compromised by adverse health conditions, such as tissue- and organ-specific perturbations of circadian rhythmicity as may be observed in shift work and jet lag, remains to be established.

Another limitation of the study is that we have had no access to the baboon blood transcriptome. This would have allowed for a comparative analysis of the extent to which tissue-specific rhythmicity overlaps in the whole blood transcriptome of these two diurnal primates.

Furthermore, rhythmicity here is described at the level of the transcriptome and how this will relate to more functionally relevant rhythmicity in the proteome is an open question.

### Outlook

Characteristics of the temporal organisation of gene expression levels in human whole blood may provide information on rhythmicity in tissues and organs. This has potential for the monitoring of tissue- and organ-specific abnormalities and effects of tissue- and organ-specific intervention. For example, monitoring of Rnf207, which is implicated in Cardiac K channel regulation, may provide new insights in the diurnal variation of Long QT syndrome and syncope. Likewise, monitoring of a set of markers for a specific organ may provide information on the timing of organ specific rhythmicity which then could be used to optimise chronotherapy for a specific organ’.

To further our insights into the validity of this approach requires a comparison of the blood transcriptome and time series of gene expression in human tissues and organs in patients with for example renal or cardiac or other abnormalities.

## Abbreviations

ADC: Adrenal Cortex

ADM: Adrenal Medulla

AMY: Amygdala

ANT: Antrum

AOR: Aorta (endothelium)

ARC: Arcuate Nucleus

ASC: Ascending Colon

AXL: Axillary Lymphonodes

BH: Benjamini and Hochberg multiplicity correction

BLA: Bladder

BOM: Bone Marrow

CEC: Cecum

CER: Cerebellum

COR: Cornea

DEC: Descending Colon

DMH: Dorsomedial Hypothalamus

DUO: Duodenum

FDR: False Discovery Rate

GO: Gene Ontology

HAB: Habenula

HEA: Heart

HIP: Hippocampus

ILE: Ileum

IRI: Iris

KIC: Kidney Cortex

KIM: Kidney Medulla

LGP: Lateral Globus Pallidus

LH: Lateral Hypothalamus

LIV: Liver

LUN: Lung

MEL: Mesenteric Lymphonodes

MGP: Medial Globus Pallidus

MMB: Mammilary Bodies

MUA: Muscle Abdominal

MUG: Muscle Gastrocnemian

OES: Oesophagus

OLB: Olfactory Bulb

OMF: Omental Fat

ONH: Optic Nerve Head

PAN: Pancreas

PIN: Pineal

PIT: Pituitary

PLSR: Partial Least Squares Regression

PON: Pons

PRA: Preoptic Area

PRC: Prefrontal Cortex

PRO: Prostate

PUT: Putamen

PVN: Paraventricular Nuclei

RET: Retina

RPE: Retinal Pigment Epithelium

SCN: Suprachiasmatic Nucleus

SKI: Skin

SMM: Smooth Muscle

SON: Supraoptic Nucleus

SPL: Spleen

STF: Stomach Fundus

SUN: Substantia Nigra

TES: Testis

THA: Thalamus

THR: Thyroid

VIC: Visual Cortex

VMH: Ventromedial hypothalamus

WAM: White Adipose Mesenteric

WAP: White Adipose Pericardial

WAR: White Adipose Perirenal

WAS: White Adipose Subcutaneous

WAT: White Adipose Tissue

## Declarations

Ethics approval and consent to participate: The protocols in which the human data were collected received a favourable opinion (EC/2007/110/FHMS & EC/2008/61/FHMS) from the University of Surrey’s Ethics committee. All participants provided written informed consent. Baboon data were collected by Mure et al. [25] and, as reported, approved by the Kenya IACUC (Institute Animal Care and Use Committee, Approval 160811) and (ISERC, (Institution Scientific and Ethical Review Committee).

## Consent for publication

Not applicable

## Availability of data and materials

The datasets analysed during the current study are available in GEO [28, 29] accession numbers GSE48113 and GSE39445.

## Author Contributions

DJD, SNA, CML, and EL conceived the analysis. CML, EL and SNA analysed the data. All authors contributed to writing the manuscript and approved to final manuscript. Acknowledgements: Not applicable

---

Supplemental Figure 1: Workflow of analyses performed. The workflow can be divided into three sections differentiated by the type of data used for the analyses: lists of rhythmic genes, gene expression levels or acrophase value. Each section describes the analyses performed and their objective(s).

Supplemental Figure 2: Overview of baboon and human orthology. Venn diagrams describing a) the number of baboon genes, as referenced by Ensembl IDs, that are orthologous to a gene in human; b) the number of baboon genes, as referenced by Ensembl IDs, that are orthologous to a human gene targeted by the microarray platform of [26]; c) the number of probes targeting genes that are both rhythmically expressed in human blood and have a baboon ortholog; d) the number of orthologous baboon/human genes that are reported as rhythmically expressed in at least one baboon tissue and human blood.

Supplemental Figure 3: Comparison of two hypothetical time series using circular cross-correlation. The time series labelled as Human (plotted in red, acrophase at 2 h) is 4 hours advanced relative to the time series labelled as Baboon (plotted in purple, acrophase at 6 h) (a). The optimal lag identified via circular cross correlation is −4 hours, indicating that the human profile is 4 hours advanced relative to the baboon profile (b).

Supplemental Figure 4: Comparison of human and baboon acrophases using circular correlation. The comparison is performed on the acrophases of the set of n genes rhythmic in both, human blood (A_H_) and in a baboon tissue/organ (A_B_). The acrophases in human blood are compared to the acrophases in the baboon tissue/organ using circular correlation r. A circular correlation and p-value are obtained for each human-baboon comparison.

Supplemental Figure 5. Prediction of the phase of rhythmicity in selected baboon tissues/organs from a set of transcripts in a human blood sample. Schematic diagram of the development and validation of a multivariate whole-blood mRNA-based predictor of tissue/organ-specific rhythmicity using a Partial Least Squares Regression (PLSR) model. Input to the model is the human blood expression X of g candidate prognostic genes in n training samples. The model’s target, Y, is the human blood sampling time relative to the acrophase of the tissue-marker gene. PLSR is used for model selection (number of latent factors and predictor genes p) via cross-validation and to fit the Ŷ =B X + B_o_ model. Validation of the model is performed by comparing the predicted vs the “observed” acrophase of the tissue-marker (blood sampling time – Ŷ) using an independent set of validation samples.

Supplemental Table 1: Overlap between orthologous human blood and baboon tissue rhythmic genes. All human/baboon orthologous genes assessed in [26] and [25]. In the columns, 1 indicates presence, 0 absence.

Supplemental Table 2: Distribution of all overlapping rhythmic genes across tissues/organs. Overlap of rhythmic genes between blood and specific tissues (red dots), and overlap of rhythmicity in tissues/organs and other tissues/organs.

Supplemental Table 3: Top ten significant gene ontology biological process and molecular function gene sets enriched within the baboon and human truly circadian genes.

Supplemental Figure 6: Effect of external phase reference marker on phase relationship in baboon and human blood transcriptome. We calculated the circular cross-correlation and associated lag between the baboon expression profile and the human diurnal expression profile for all overlapping rhythmic genes. The circular cross-correlation and lags were calculated in every baboon tissue/organ based on alignment with start of light, midpoint of light, and end of light. The resulting three lags per gene, per baboon tissue, were ranked such that the alignment with the smallest lag is given the top rank (1). a) Heatmap and b) histogram of ranked lags of all overlapping rhythmic genes, where ranking was performed within gene across the three different alignments.

Supplemental Table 4: Circular cross-correlation lags for the clock genes that were rhythmic in human blood and identified as rhythmic in the greatest number of baboon tissues/organs.

Supplemental Table 5: Clustering results of circular clustering analysis of all overlapping tissue/organ-specific rhythmic genes in human blood and baboon tissues/organs combined.

Supplemental Figure 7: Circular clustering analysis was used to characterise the distribution of tissue/organ-specific rhythmic genes in human blood and baboon tissues/organs. Clustering of acrophases of all overlapping tissue/organ-specific rhythmic genes in human blood and baboon tissues/organs combined. The number of clusters was limited to two. In all panels acrophases are coloured by cluster membership and are expressed as hours since lights on. a) Scatter plot of acrophases of human and baboon genes; b) distribution of acrophases in human blood; c) distribution of acrophases in baboon tissues/organs.

Supplemental Figure 8: Summary of alignment of rhythmic genes in baboon tissues/organs and their human rhythmic orthologs in blood. a) Circular correlations of acrophases. b) Circular cross correlation of gene expression temporal profiles. c) Mean lag in hours between human blood and baboon tissues/organs expression profiles. d) Standard deviation of the lag. Negative lags indicate that the timing of a human rhythm precedes the baboon rhythm. Asterisks indicate Benjamini-Hochberg corrected significance of circular correlation * P < 0.05; **P < 0.01; ***P < 0.001.

Supplemental Table 6. Biomarker candidate genes. Sets of probes targeting orthologous genes rhythmic in both human blood and the selected baboon tissues/organs. Column F marks (with 0) genes excluded from the sets. Column G indicates (with 1) genes used in the 1-sample PLSR model. Column H indicates (with 1) genes used in the 2-sample PLSR model.

Supplemental Figure 9: Blood based biomarkers for tissue/organ-specific rhythmicity: performance when sleeping out of phase. Performance of one-and two-sample models derived from the training set when used to predict timing of ANT and PVN rhythmicity in the sleeping out of phase condition validation set. Predicted timing of ANT vs. sample clock time for each sample in the validation set for a) the one-sample and b) two-sample PLSR model. c) Proportion of predictions of timing of ANT vs. cumulative absolute error for the one- and two-sample PLSR model. Predicted timing of PVN vs. sample clock time for each sample in the validation set for d) the one-sample and e) two-sample PLSR model. f) Proportion of predictions of timing of PVN vs. cumulative absolute error for the one- and two-sample PLSR model. The cumulative frequency distribution of absolute error can be used to identify the proportion of samples that can be predicted within a given error range, where the highest proportion combined with smallest error range denotes the better prediction model.

