## Supplemental Figure 1 for "Overlap of expression and alignment of diurnal and circadian rhythmicity in the human blood transcriptome with organ and tissue specific rhythmicity in a non-human primate"

| Input | Analysis | Objectives |
| --- | --- | --- |

### List of rhythmic genes

- Human blood
- Baboon tissues/organs

**Set operations**

- Overlap of rhythmic genes
- Overlap of tissue-specific rhythmic genes

**Enrichment analysis**

- Characterise overlap and distribution of all overlapping rhythmic genes in human blood and across baboon tissues/organs
- Detect genes that are rhythmic in only one baboon tissue/organ in the rhythmic transcriptome in human blood
- Compare diurnal vs circadian overlap of all overlapping rhythmic genes and of tissue-specific overlapping rhythmic genes
- Identify significantly enriched GO terms in the overlapping rhythmic gene sets

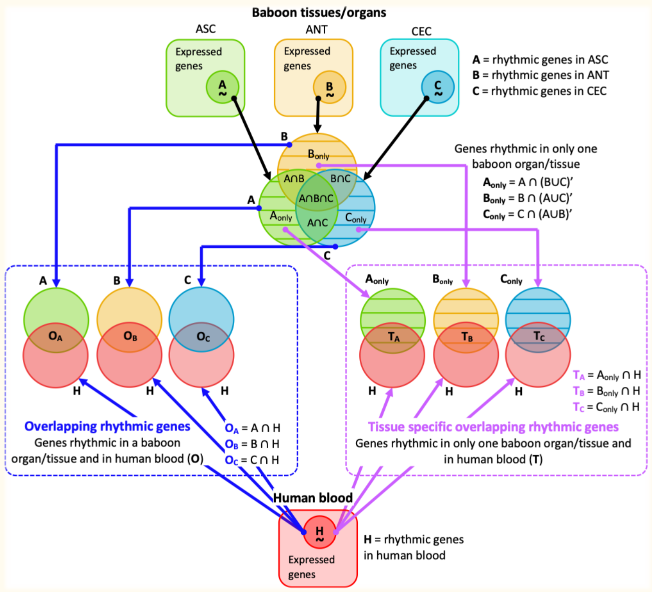


### Gene expression levels

- Human blood
- Baboon tissues/organs

**Mean expression level comparison**

**Selection of reference alignment**

**Circular cross-correlation analysis**

- Explore relationship between average level of expression and the detection of rhythmicity
- Identify which phase reference point yields the closest alignment between the baboon and human transcriptomes
- Characterise alignment of core clock genes under diurnal conditions

### Acrophases

- Human blood
- Baboon tissues/organs

**Circular correlation analysis**

**Acrophase difference analysis**

**Circular clustering analysis**

**Enrichment analysis**

- Characterise alignment of core clock genes under diurnal conditions
- Compare timing of all overlapping rhythmic genes in the baboon and human blood, separately for each tissue
- Identify whether the timing of tissue-specific overlapping rhythmic transcripts is similar to their timing in human blood
- Characterise the distribution of overlapping rhythmic genes within the bimodal peak distribution for human blood and baboon tissues/organs
- Identified significantly enriched GO terms in the gene sets clustered by acrophase values
