## Supplementary figures and images for "Overlap of expression and alignment of diurnal and circadian rhythmicity in the human blood transcriptome with organ and tissue specific rhythmicity in a non-human primate"

### Supplemental Figure 2

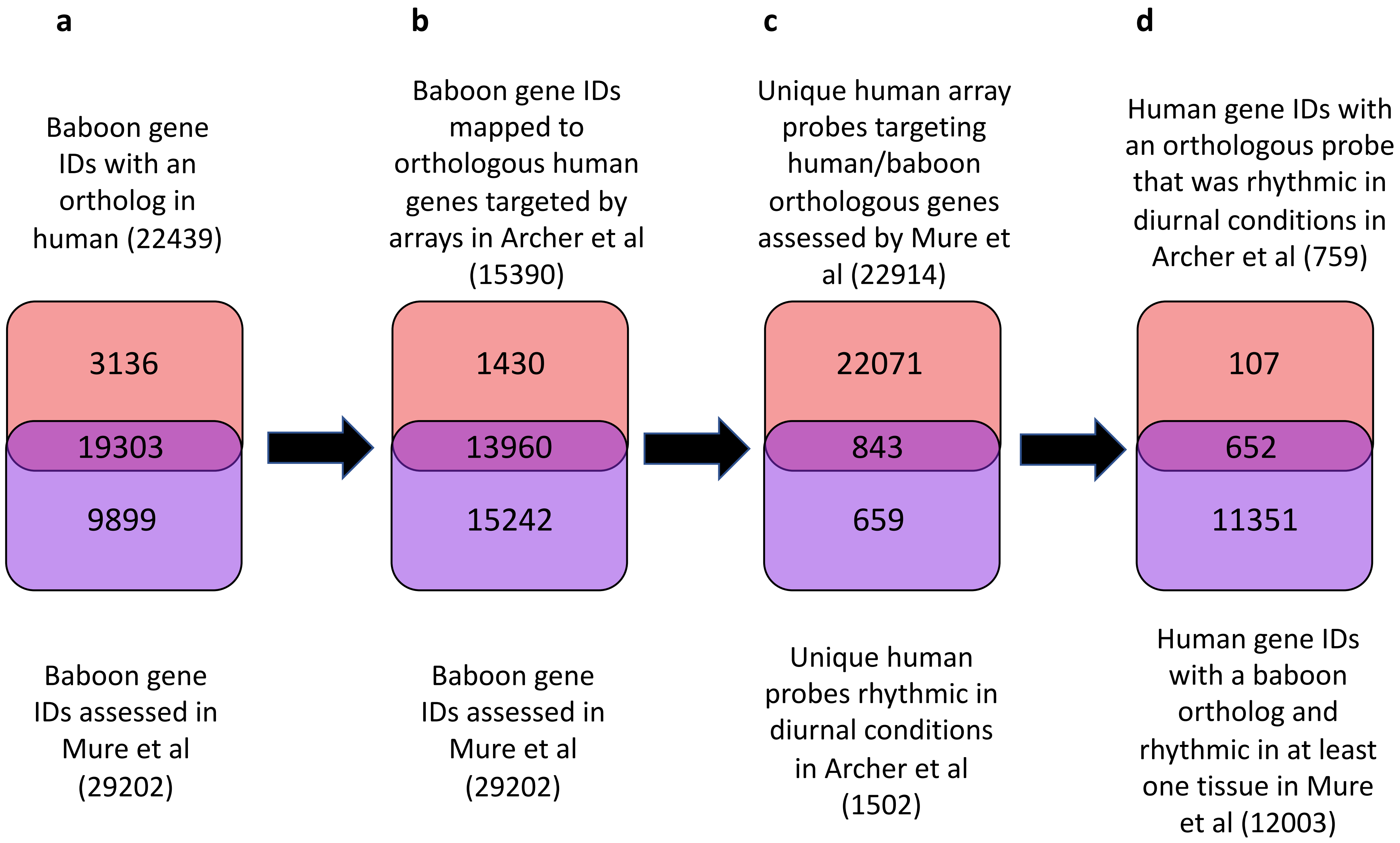

### Supplemental Figure 3

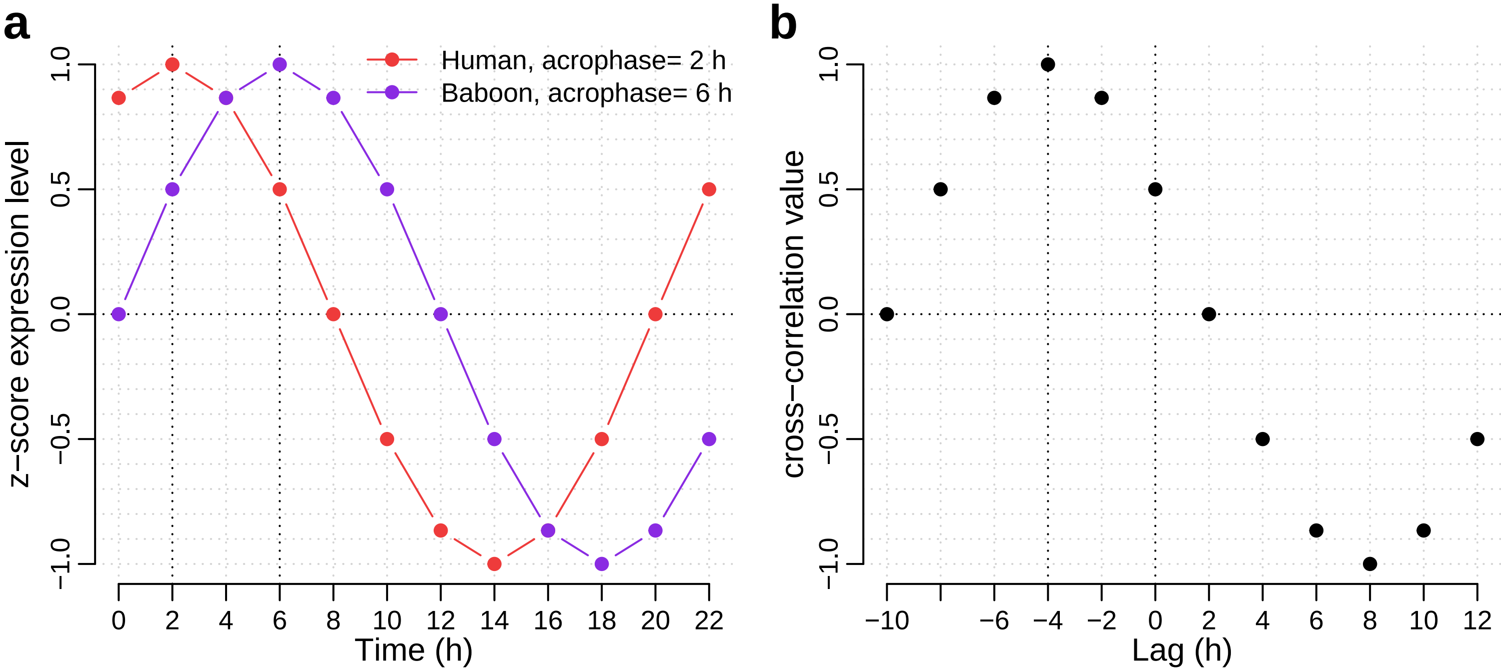

### Supplemental Figure 4

**Comparison of baboon and human acrophases (circular correlation)**

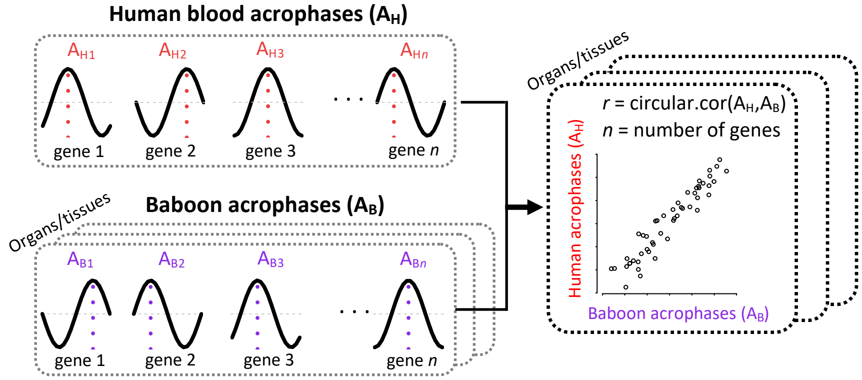

### Supplemental Figure 5

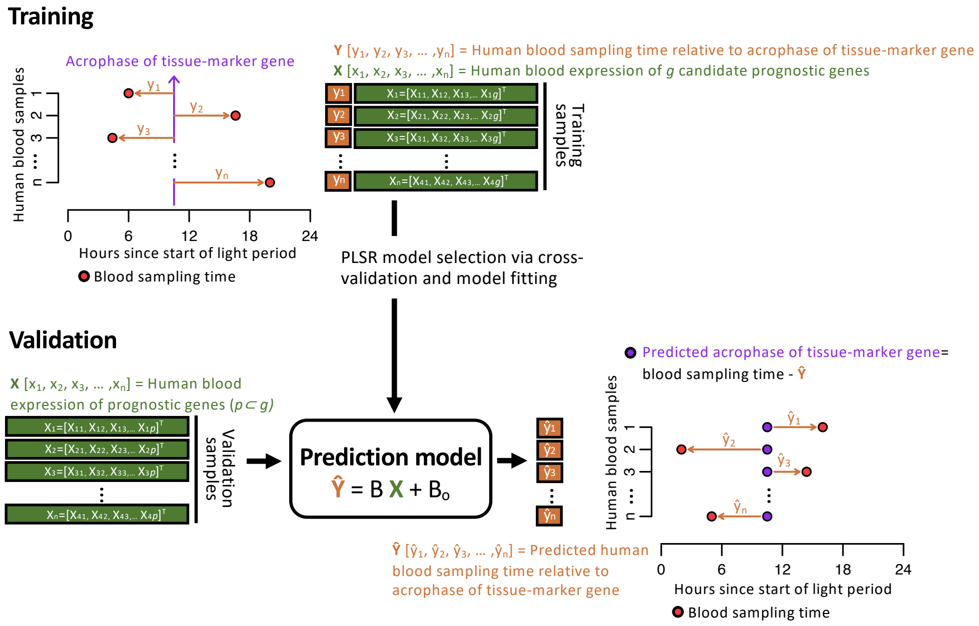

### Supplemental Figure 6

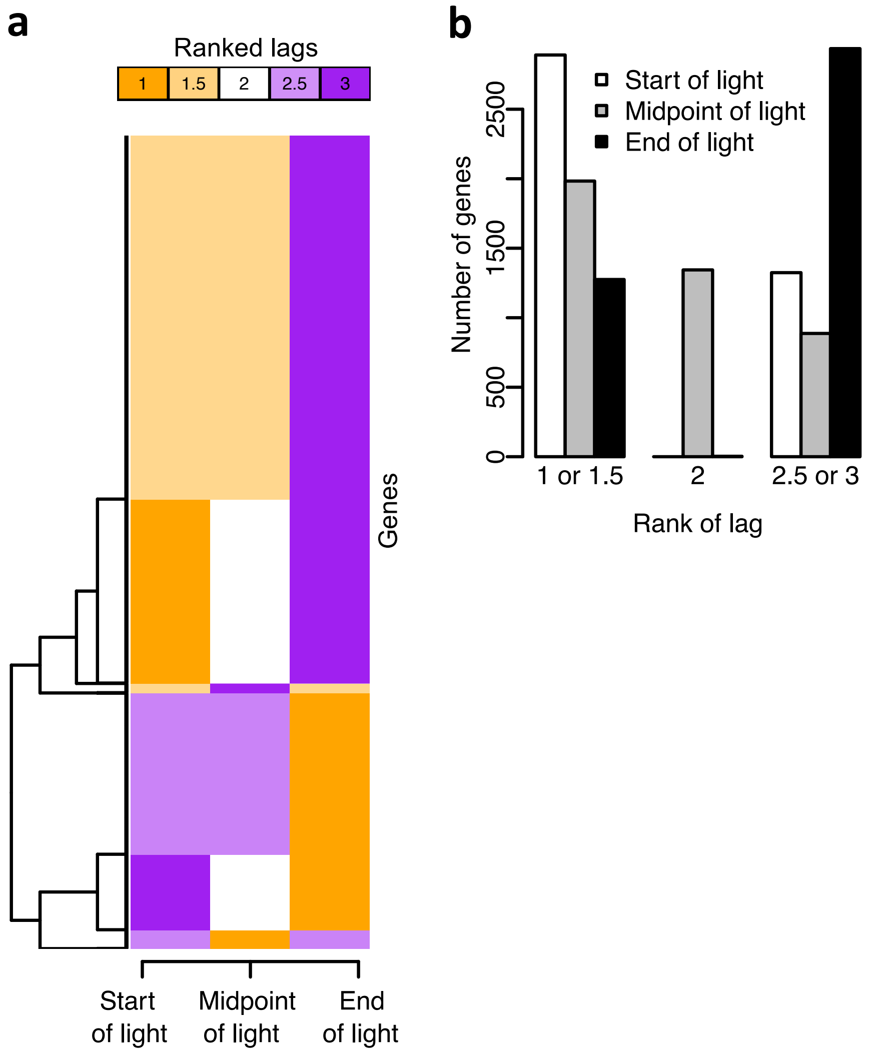

### Supplemental Figure 7

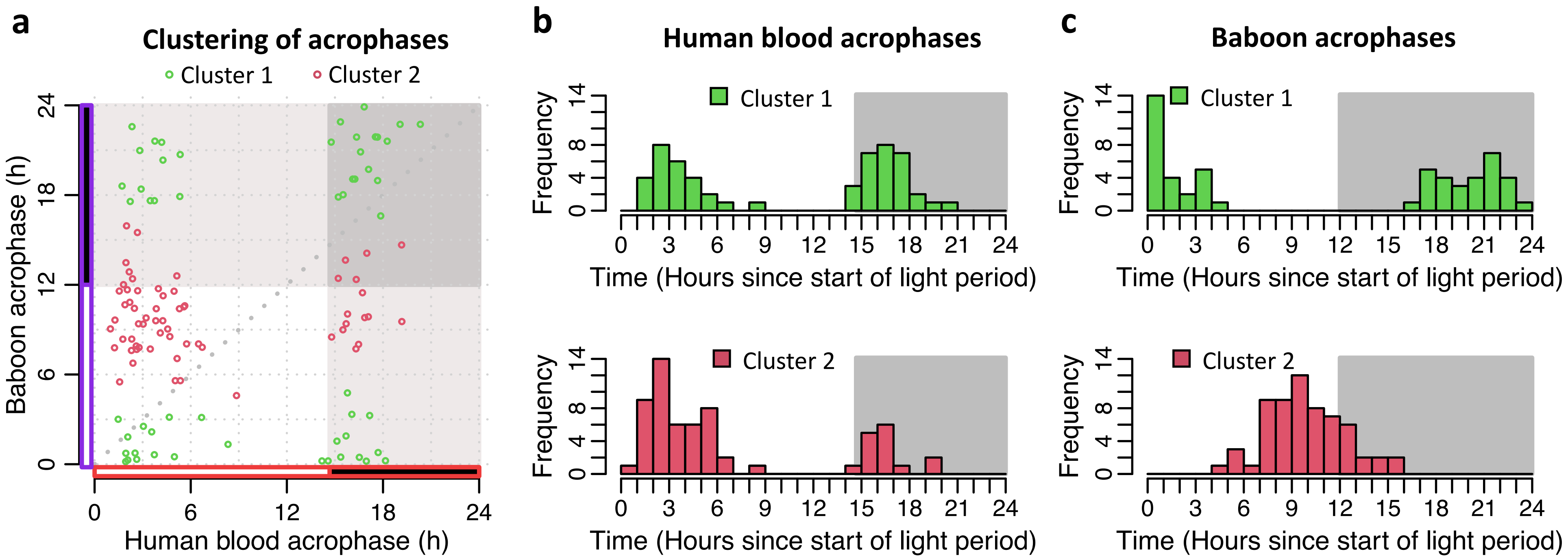

### Supplemental Figure 8

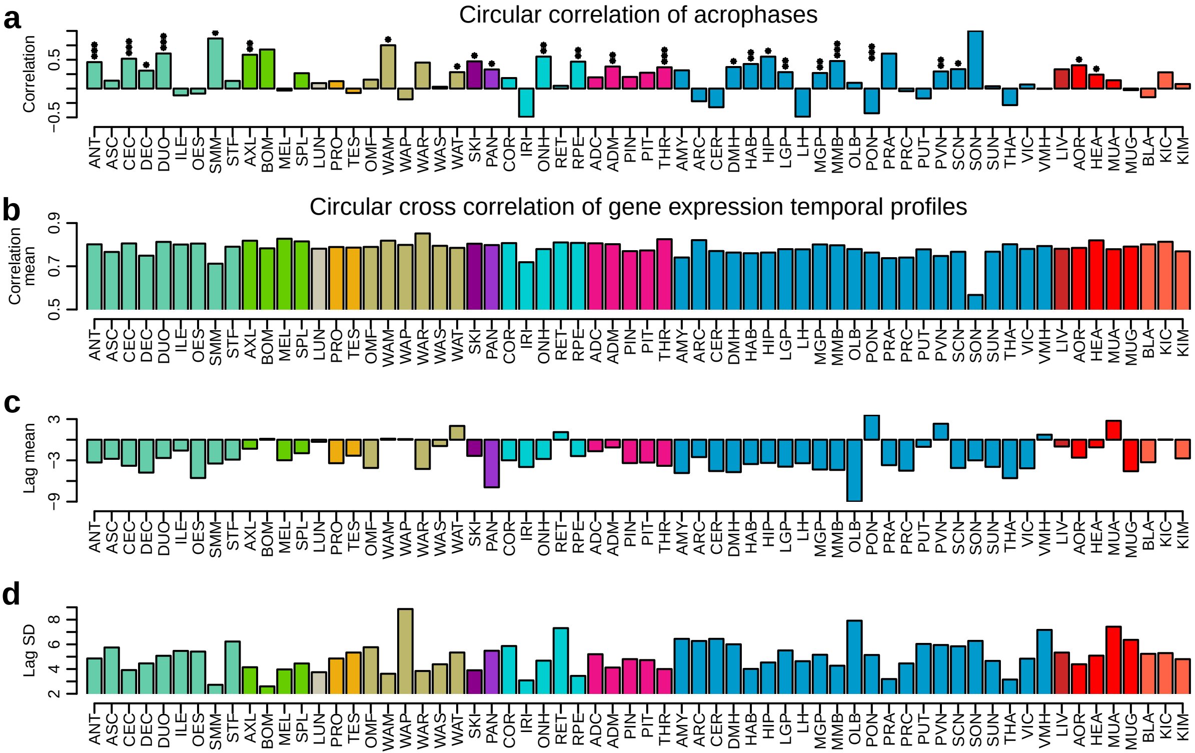

### Supplemental Figure 9

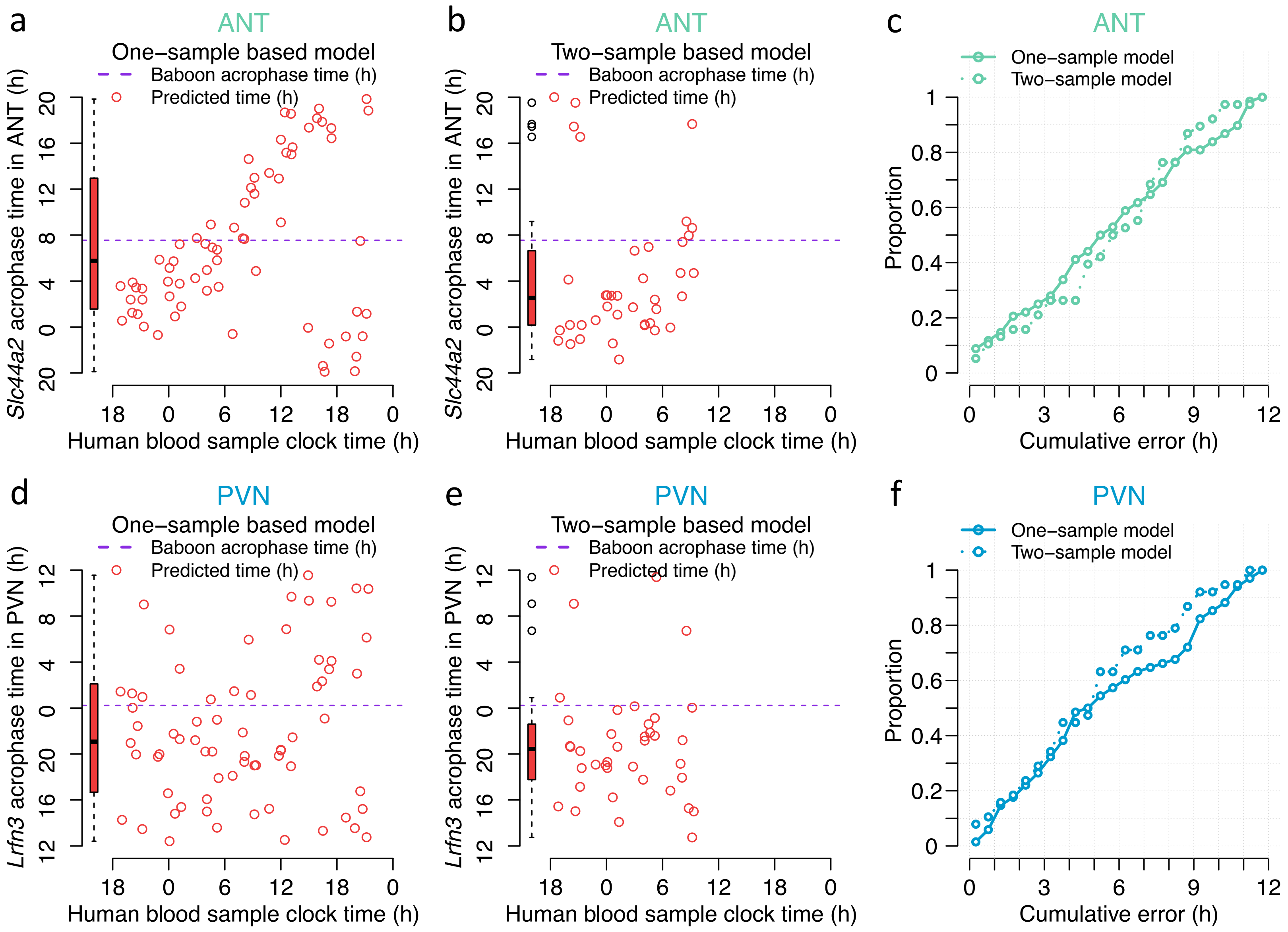
