## Supplemental Table 3 for "Overlap of expression and alignment of diurnal and circadian rhythmicity in the human blood transcriptome with organ and tissue specific rhythmicity in a non-human primate"

| Gene Set | Description | P value | FDR |
| --- | --- | --- | --- |
| GO:0001818 | Negative regulation of cytokine production | 0.00024917 | 0.28206 |
| GO:0042092 | Type 2 immune response | 0.0034364 | 1 |
| GO:0044262 | Cellular carbohydrate metabolic process | 0.0063316 | 1 |
| GO:0032613 | Interleukin-10 production | 0.0068721 | 1 |
| GO:0043177 | Organic acid binding | 0.0072210 | 1 |
| GO:0022613 | Ribonucleoprotein complex biogenesis | 0.0084140 | 1 |
| GO:0070206 | Protein trimerization | 0.0096451 | 1 |
| GO:0003707 | Steroid hormone receptor activity | 0.010162 | 1 |
| GO:0001562 | Response to protozoan | 0.011037 | 1 |
| GO:0042110 | T cell activation | 0.011243 | 1 |

Supplemental Table 3. Top ten significant gene ontology biological process and molecular function gene sets enriched within the baboon and human truly circadian genes.
