## Supplemental Text for "Overlap of expression and alignment of diurnal and circadian rhythmicity in the human blood transcriptome with organ and tissue specific rhythmicity in a non-human primate"

**S1) Over/under-representation analysis of known blood immune-cell type gene markers of the overlapping rhythmic genes**

Over and under representation of blood immune-cells, as indicated by immune cell markers defined in [42], within the day or night cluster (see main text), of overlapping rhythmic genes identified in a given tissue/organ was assessed using a two-sided Fisher exact test. Here, overlapping genes within a given cluster, within a tissue/organ, were directly compared with each set of orthologous immune cell markers. Supplemental Table 7 provides the observed vs expected values, and associated p values, for each cluster, tissue and immune cell type combination.

**S2) Functional annotation of the genes that predict the phase of the PVN and Antrum**

Functional annotation of the genes that predict the phase of the PVN and antrum showed that the the predictors were related to cell cycle/cell growth/apoptosis (e.g. *Tspan4*, *Hpn*, *Aatk*, *Klhdc8B* in the PVN; *Hipk2*, *Ccnh*, *Ralb*, *Ranbp1*, *Rps27a* in the antrum), ubiquitination/protein degradation/autophagy (*Hectd2*, *Atg10*, *Dcun1d1*, *Birc2*, *Atg7* in the PVN; *Ulk1*, *Wdfy3*, *Cnot4*, *Erlec1*, *Kctd21*, *Trim25*, *Trim50* in the antrum), and transcription/translation (*Aarsd1*, *Tcea3*, *Trmt61a*, *Znf639*, *Zfp62*, *Hsf2*, *Wdr74*, *Larp7*, *Eef1g*, *Rps5* in the PVN; *Phf21a*, *Pus7*, *Gnl2*, *Znf395*, *Snrpd1*, *Exosc8*, *Rpl17* in the antrum).

**S3) Effect of Baboon’s sampling scheme on the identification of rhythmic genes and their overlap with human blood rhythmic genes.**

The baboon dataset consists of a single-replicate time-series sampled every two hours for 22 hours, producing 12 samples within the same 24 hours cycle. A 13^th^ time point would be required to have at least one pair of samples 24 hours apart.

We explored the impact of 12 vs 13 samples on the percentage of correct classifications by using Time trial <https://www.biorxiv.org/content/10.1101/2020.04.15.043695v2> and CircaInSilico (<https://5c077.shinyapps.io/Circa_in_Silico/>). In both cases 12 samples gives a percentage correct classification which is very similar to 13 samples.

*CircaInSilico*

We used the tool CircaInSilico [Hughes et al. JBR 2017] to generate a synthetic rhythmic mRNA transcript dataset that follows the Baboon sampling scheme and a second dataset with an equivalent sampling scheme using 13 samples instead of 12. We used MetaCycle to identify the number of rhythmic genes in the synthetic datasets and calculated the number of false positives and false negatives:

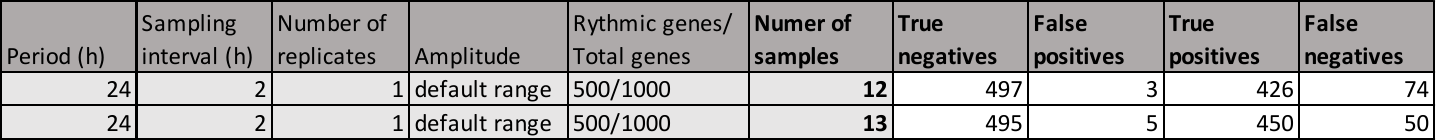

We observed that the number of false positives was not affected by the use of 13 vs 12 samples. In both cases the number of false positives is extremely low (3 for the 12-sample dataset and 5 for the 13-sample dataset). However, there was an increase in false negatives when using 12 samples relative to 13 samples: 85% of rhythmic genes are identified in the 12-sample dataset, while 90% of rhythmic genes are identified in the 13-sample dataset.

*TimeTrial*

We used the tool TimeTrial [Ness-Cohn 2020] to evaluate the effect of the Baboon sampling scheme in the identification of rhythmic genes. Noise levels correspond to additive Gaussian noise as a percentage of the wave form amplitude. We compared the Baboon sampling scheme, with 12 samples, with an equivalent sampling scheme using 13 samples:

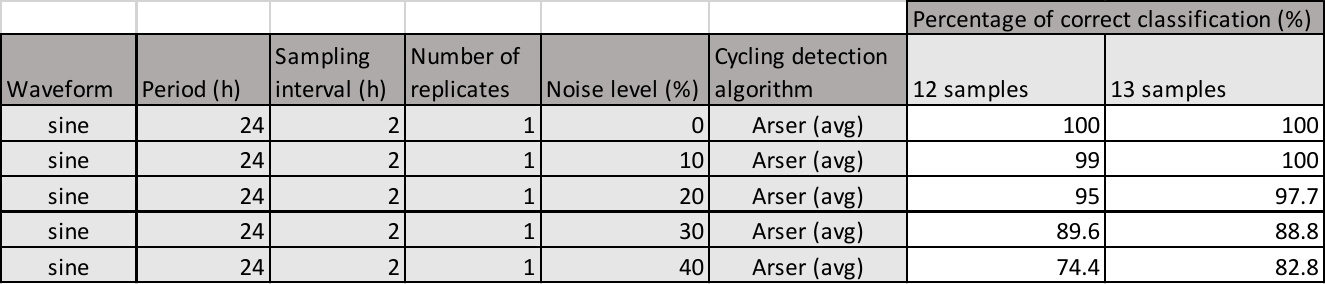

Please note that we had to adapt this tool to conduct these simulations. Code is available upon request.

*Significance of rhythmic sets overlap*

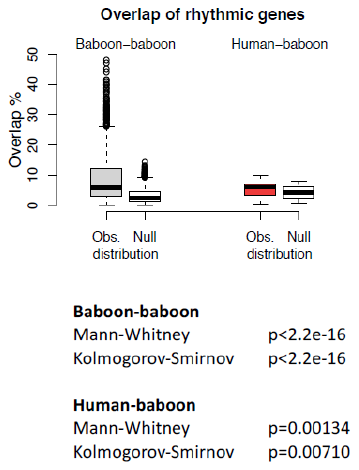
We also conducted an analysis of the likelihood that the overlap between the sets deviated significantly for the overlap between two random sets. To generate the null distribution of overlaps we randomly shuffled the Baboon rhythmicity results table in two ways (each 50 times) first by keeping the same number of rhythmic genes in the table and then by keeping the same number of rhythmic genes within each baboon tissue/organ. We calculated the overlap of the shuffled tables with the set of human rhythmic genes and compared the distribution of observed overlaps with the null distribution using the Mann-Whitney and Kolmogorov-Smirnov tests. We found that the observed distribution of overlaps has significantly higher values (p<0.05) than what is expected from the overlap of two random sets.
